# Activity and specificity trade-offs in adenine base editors

**DOI:** 10.64898/2026.07.20.739671

**Authors:** Maria Lukarska, Luke M. Oltrogge, Hunter Nisonoff, Yaeko Long, Cynthia I. Terrace, Akosua Busia, Cristal Aquino, Shin Kim, Jennifer Listgarten, David F. Savage

**Affiliations:** Department of Molecular and Cell Biology, University of California, Berkeley, Berkeley, 94720, CA, USA; Innovative Genomics Institute, University of California, Berkeley, Berkeley, 94720, CA, USA; Howard Hughes Medical Institute, University of California, Berkeley, Berkeley, 94720, CA, USA; Department of Electrical Engineering & Computer Sciences, University of California, Berkeley, Berkeley, 94720, CA, USA; Center for Computational Biology, University of California, Berkeley, Berkeley, 94720, CA, USA

**Author notes:** These authors contributed equally to this work (Maria Lukarska, Luke M. Oltrogge, Hunter Nisonoff). Denotes co-corresponding authors.

## Abstract

Adenine base editors (ABEs) are CRISPR effectors that introduce A-T to G-C transitions in the genome using a nucleotide deaminase fused to a Cas protein. ABEs have been evolved to have very high editing efficiency, but off-target editing effects compromise their precision and hinder their applications. Here we explore the activity and specificity relationship of ABEs using a combination of machine learning-guided design and high-throughput screening. We designed a diverse library of 12,000 variants and built quantitative bacterial selection systems that allowed us to simultaneously measure their on-target and off-target editing. We found that the ABEs were fully described by the single dimension of intrinsic deaminase activity with no evidence for independent specialization with respect to local sequence context, editing window width, RNA editing, or genotoxicity. These results were supported by *in vitro* studies and consistent with editing experiments in mammalian cells. Finally, the activity and specificity trade-offs were recapitulated among previously reported engineered variants and a selection of library variants spanning the activity spectrum. Our results suggest that fundamental architectural improvements will be necessary to transcend the activity and specificity limitations for the next generation of ABEs.

## Introduction

Precise and targeted editing of the genome has the potential to correct pathogenic mutations in humans or engineer desirable traits in crops. Base editors (BE) are genome editing tools that can make single nucleotide changes to DNA. Adenine base editors (ABEs) comprise an adenine deaminase domain fused to a catalytically impaired CRISPR-Cas protein. ABEs can deaminate an adenine (A) to inosine (I) within the R-loop formed by Cas9. The inosine is recognized as a guanosine (G) during replication, leading to an A to G transition. ABEs were engineered from TadA, a tRNA deaminase which edits the anticodon loop of tRNAs, over several directed evolution campaigns^1,2^, leading to the highly active editor, ABE8e^2^.

ABEs have been successfully used in different clinical models and human patients, however, concerns remain regarding specificity and precision^3–11^. ABEs edit a target R-loop formed by Cas9 within an editing window of around 10 nucleotides. They can also edit at non-cognate sites, leading to off-target edits. The off-target editing of ABEs can be classified in several ways including: bystander edits, Cas-dependent and Cas-independent DNA off-targets, and transcriptome-wide RNA off-targets^12^. Specifically, bystander edits refer to editing by ABEs of additional A’s present in the non-target strand window aside from the A’s intended to be edited. Cas-dependent off-targets are due to transient R-loop formation at non-cognate sites that contain mismatches to the gRNA allowing ABEs with high intrinsic activities to edit this R-loop. Cas-independent off-targets are caused by the ABEs editing random regions of DNA that likely become accessible during cellular processes like transcription or replication. Finally, ABEs can edit single-stranded RNA (ssRNA), leading to transcriptome-wide edits^13–15^.

ABEs have been shown to induce geno- and cytotoxicity from genome- and transcriptome-wide off-target effects prompting multiple efforts to develop specific deaminase variants^16,17^. Different strategies to increase specificity include rational structure-based design^13,15,18–21^, continuous evolution methods^22,23^ and reverting mutations from the directed evolution campaigns^24^. Some engineered variants have been used clinically, including ABE8-V106W which was administered to an infant to correct a severe urea cycle mutation in the liver^9^. A challenge in the field remains to engineer highly precise and active ABE variants with reduced risk of genotoxicity.

Recent developments in machine learning (ML)-guided design have enabled the exploration of protein sequence space^25^. An advantage of these approaches is that a single round of designs can sample a large breadth of sequence diversity while maintaining function. Classically, this would require multiple rounds of experimental diversification such as directed evolution. Here, we developed a high-throughput bacterial screen that allowed quantitative assay of thousands of ML-designed ABE variants across different edit targets and conditions. We discover a tight trade-off between specificity and activity over our diverse library and previously published variants. This trade-off is recapitulated in human cells among a selection of our library variants spanning a wide activity spectrum. Together, these results suggest that previous engineering campaigns on the ABE deaminase domain have optimized along the single dimension of activity without any change to specificity.

## Results

### Library design and assays for measuring on-target activity

As a first step in understanding the activity and specificity of base editors we wanted to test if we could increase the activity of the current state-of-the-art ABE8e (hereafter referred to as ABE8) using a ML-based approach to library design. Specifically, we created a diverse library of ABE variants using an ML model comprising two components, each with a different information source (Figure 1A). The first component modeled protein family-based evolutionary information. The second component modeled ABE8 structural information via an inverse folding model. Each component of the model was trained separately before being combined into a single model. The evolutionary model, trained on a multiple-sequence alignment of natural TadA homologs, captures the sequence constraints and amino-acid covariation imposed by evolution; the inverse-folding model captures how likely a candidate sequence is predicted to fold into the ABE8 backbone structure. The overall model averages the scores (i.e., the log likelihood) of these two models (Figure S1A). To use the overall model for design, we defined a library objective function that trades off high average model score, with library diversity and then optimized this objective function to yield a designed library (see Methods, Figure S1B).

**Figure 1.**
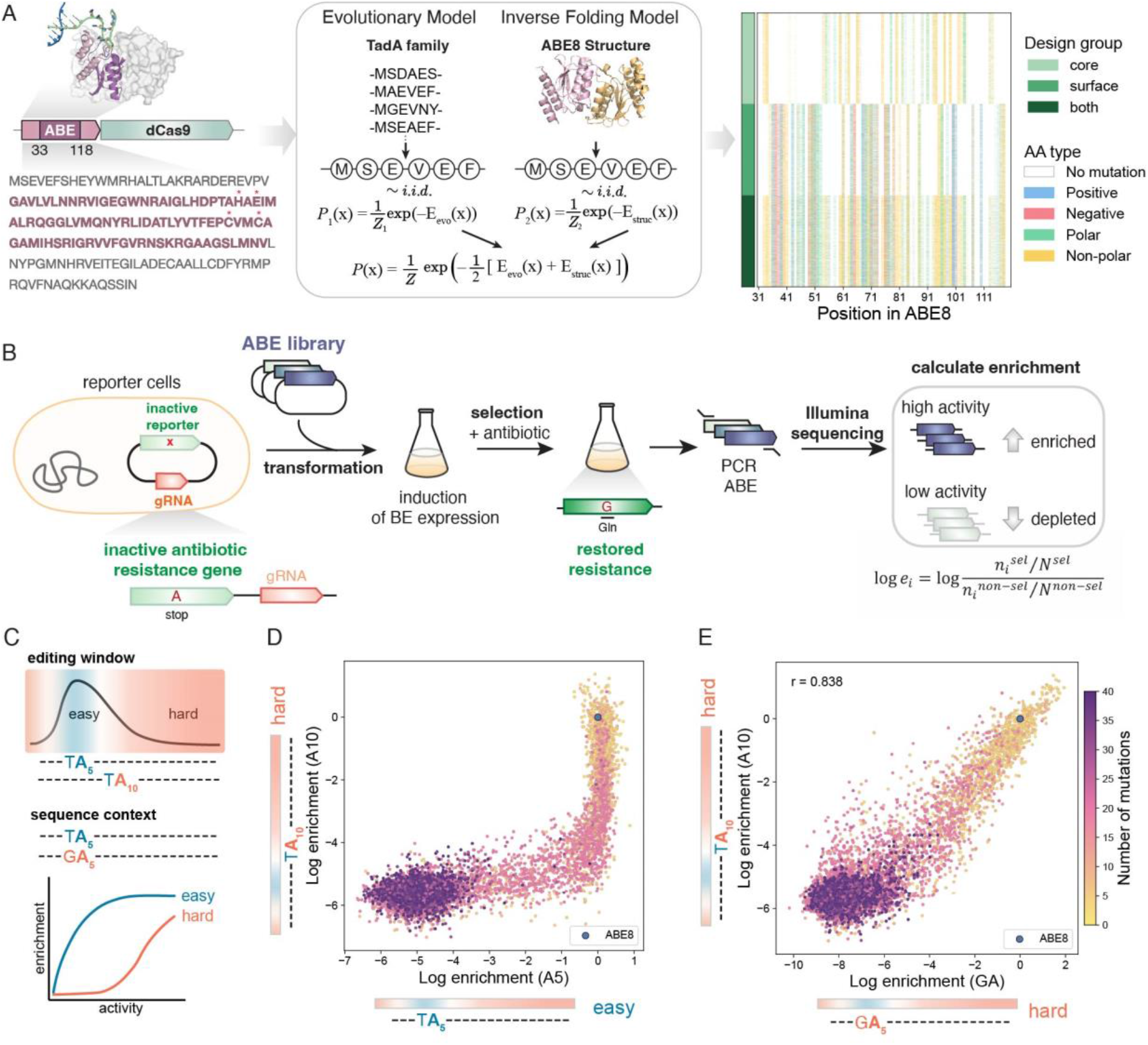
ML-guided design and high-throughput selections of diverse ABE libraries. A.Left: highlight of the ABE region that is diversified is shown in purple, below: sequence of the region with red asterisks indicating active site residues; above: structure of the ABE dimer within the ABE8-Cas9 complex shown in surface representation, one of the monomers shown in cartoon, diversified region in purple, non-target DNA in green and target DNA in blue (PDB: 6VPC). Middle panel: combination of two ML models used for generating the library, evolutionary and inverse folding model. Right: illustration of the library diversity, variants are grouped by design group (core, surface, both core and surface), indicated in shades of green, each type of substitution is colored by the type of mutation (positive, negative, polar or non-polar). Each variant is a separate row. B. Schematic of the on-target selection assay. *E*.*coli* cells are pre-transformed with a reporter plasmid carrying the inactivated antibiotic resistance gene and a guide RNA targeting it. The library is transformed, ABE expression induced and cells are grown under selective and non-selective conditions. The diversified region is sequenced and the enrichment of variants is calculated from the log ratio of counts under selective over non-selective conditions. C. Two types A. of assays are used to capture different parts of the library’s activity dynamic range: the easy selections (with a target in the middle of the editing window, A5, and target A preceded by a pyrimidine, TA) allow to resolve the lower activities, and the hard selections (which can be hard by virtue of the target A being at the end of the editing window, A10, or in the middle of the window, but preceded by a G, GA) allow to measure the highly active variants. D. Examples of an easy and hard assay capturing different parts of the dynamic range. Log enrichments of the variants in one example of an easy selection and one of the hard selection. Variants are colored by their number of mutations compared to ABE8. ABE8 is indicated in blue. E. Correlation between log enrichments in the two hard assays (A10 and GA). Variants are colored by the number of mutations. Replicate Pearson correlation (r) is shown on the upper left.

We focused on a contiguous region of the ABE that comprises the active site and most of the core structure (region 33-118) for purposes of oligo synthesis costs, cloning, and sequencing simplicity (Figure 1A, left). Additionally, we were aiming to generate variants with similar or higher activity than ABE8, hence we fixed the active site residues, as well as other positions that have been shown to be important for high editing activity in previous evolution campaigns leading to ABE8^1,2^ (Table S1). The rest of the protein was kept identical to ABE8.

To explore different regions of the protein, we ran the library design procedure with three different strategies mutating: i) buried (core) positions, ii) exposed (surface) positions, and iii) both surface and core simultaneously. For the core-only and surface-only strategies we allowed up to 20 mutations from the ABE8 parent, while for the combined core-and-surface strategy we allowed up to 40 (Figure S1B). Pooling the three strategies yielded a final library of 12,000 designs spanning a wide range of mutational loads (up to 40 substitutions from ABE8) (Figure 1A, right).

Next, we wanted to characterize the activities of the designed library. Antibiotic reversion has been widely used for bacterial screening: an antibiotic selection marker is inactivated by a single mutation that confers functional resistance upon base editing ^1,2,24,26^. We leveraged this principle to create a quantitative bacterial selection where cells carrying base editors survive antibiotic challenge with a fitness that depends on the editing level (Figure 1B). Direct sequencing and counting of the variants in selective and non-selective conditions can be used to calculate an enrichment value, which reflects the relative activity of the base editor variants in the library. We fused our ABE library to SpRY-Cas9, an engineered PAM-broadened variant of SpCas9, which enabled more flexible design of selection targets.

To test for target sequence bias, we designed several selection reporters that required targeting a variety of spacers in the antibiotic resistance gene (Table S2, Supplementary sequences). Assay of the library in different selection conditions revealed that each target sequence had a different “difficulty” of editing, allowing capture of the range of activities inherent to the library (Figure 1C). When the selections were based on an edit at the peak of the editing window (hereafter “easy” selections, i.e. A5 in the protospacer), we could differentiate activities in the lower and middle parts of the dynamic range, but the activities at the high end would saturate and be indistinguishable (Figure 1D). Since we were interested in variants that have higher activity than ABE8, we designed two additional screens based on “hard” targets, where the required edit is at the edge of the optimal window (A10 selection) or followed by a G (GA selection). The latter’s difficulty is due to the inherent preference for a preceding pyrimidine, inherited from the TadA ancestor^1^. The two types of hard selections allowed distinguishing the high activity variants from one another. Moreover, these assays correlated with each other, confirming that both assays are measuring base editor activity and not any substrate preference or other bias (Figure 1E). We focused on the hard selections, as they covered the higher end of the range of activities.

The activity of the variants was, on average, inversely related with the mutation count, wherein variants closer in sequence space to ABE8 tended toward higher activity while increasing the number of mutations generally lowered the activity (Figure 1D,E). However, we still observed variants carrying up to 20 mutations with activities comparable to ABE8. This approach allows us to sample a significant sequence and functional diversity that is difficult to access through experimental evolution methods. We obtained thousands of diverse variants that have detectable activity within the full mutation range (1-40). The library yielded 170 variants with higher activity than ABE8 based on the mean enrichment in the two hard assays (Figure S2). The strong correlations between biological replicates (Figure S3A), as well as between different assays relying on distinct spacer sequences (Figure 1D), show that the bacterial selections robustly measure base editing activity of large variant libraries. To confirm that the readout was not specific to the SpRY-Cas9 variant, we also cloned the library and tested it with SpCas9 (Figure S3B). Even though the selections were more qualitative, they followed the same general shape and pattern of inverse correlation with mutation number.

### Calibrating library enrichment values with intrinsic deamination activity

Next, we selected a subset of ten variants that span the dynamic range of our library to thoroughly characterize across experimental modalities (Figure 2A). We hypothesized that the difference in activities measured via high-throughput assay are independent of the Cas protein and reflect the enzymatic deamination rates of the variants. To test this, we recombinantly expressed and purified the deaminase domains of selected ABE variants. Eight of the variants were overexpressed, soluble, and could be purified to homogeneity for *in vitro* assays (Figure S4).

**Figure 2.**
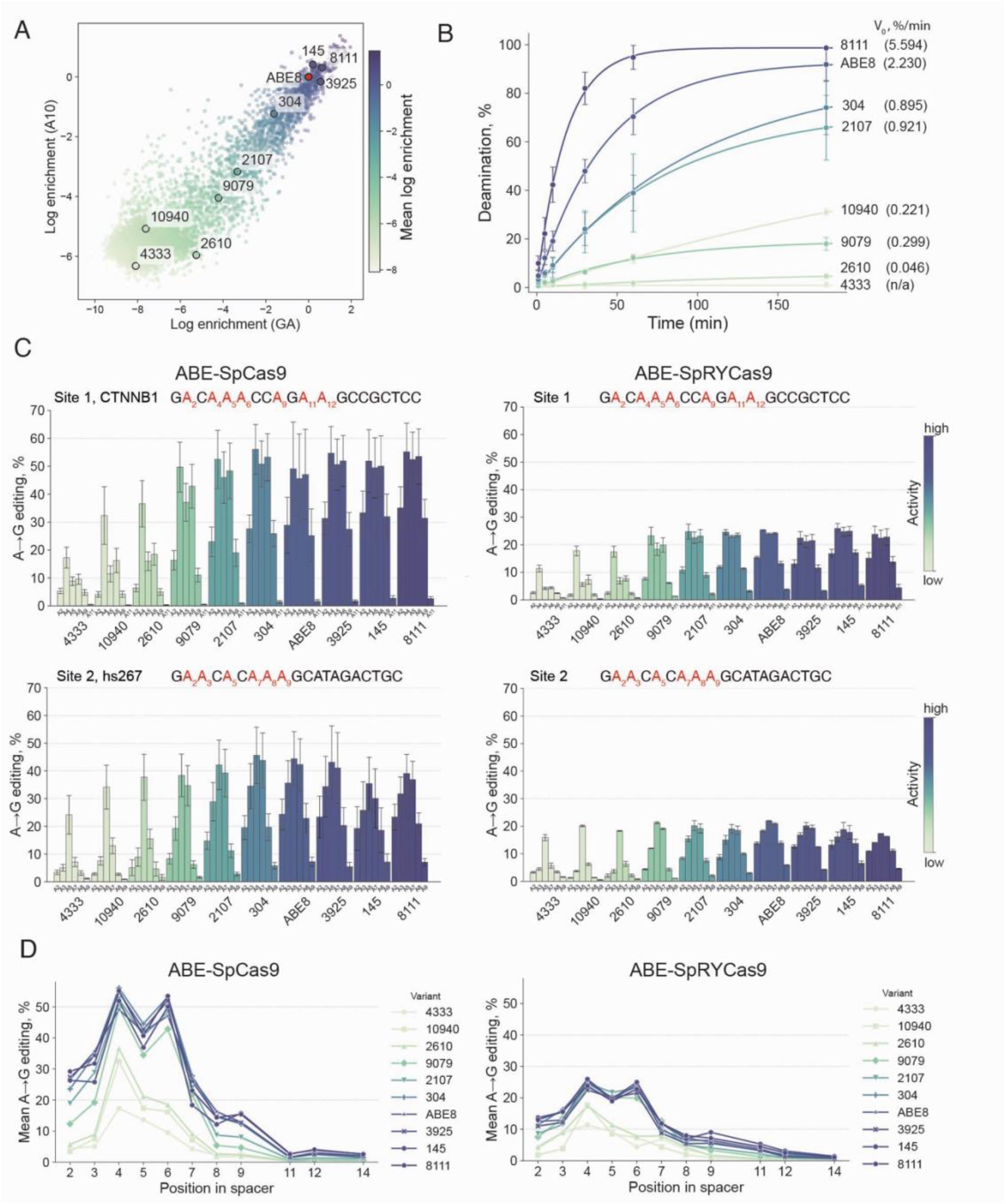
On-target activity of selected library variants. A. A subset of variants spanning the activity range and chosen for further characterization are highlighted on the correlation plot of the two hard assays. Variants are colored by their mean log enrichment in these two assays and this color scheme is used to color-code the variants on all subsequent plots. B. Deamination rates of selected variants measured by direct inosine detection in a nanopore-based assay. *In vitro* activity of the variants increases as their activity measured in the selection assays increases. Variants are colored as shown on panel A and are indicated by their number in the designed library. Deamination rates are shown on the right (V_0_). C. On-target editing on two endogenous HEK293T loci with the ABE variants fused to either nSpCas9 (left panels) or nSpRY-Cas9 (right). Edited positions shown on the bar graphs are highlighted in red in the sequence. Bars represent mean editing of three biological replicates, each averaged from three technical replicates. Error bars show standard deviation of the biological replicates. D. Editing window representing mean normalized editing at each position across all tested sites (n = 4) for the SpCas9 and SpRY-Cas9-fused ABE variants.

In order to measure the deamination activity of our selected variants, we established a direct assay that measures A to I conversion using nanopore sequencing (Figure S5, Figure 2B). We leveraged the ability of nanopore-based sequencing to directly read the inosines in a target DNA, which facilitates multiplexing of biochemical assays (see Methods). Briefly, the ABE deamination activity of the variants was tested on a single-stranded synthetic substrate, mimicking the single stranded non-target DNA of the R-loop. Upon quenching, a complementary strand was added to form a double stranded DNA, and different barcodes were ligated at each time point. The samples were then sequenced, directly detecting the inosine produced by the deaminases, leading to a robust and quantitative readout of ABE enzymatic activity.

Similarly to previous reports^26,27^ we were able to detect activity in single-turnover conditions at high enzyme concentrations. As shown previously for ABE-Cas9 fusions, we also found that ABE proceeds with limited catalytic turnover. This precluded measuring Michaelis-Menten kinetics and deriving standard *k*_*cat*_ and K_M_ values, so we instead measured the activity under single-turnover conditions and derived V_0_ as an overall deamination rate. Also consistent with previous assays using other ABE variants^27^, we observed different final amplitudes of the reactions at long time points. Our data shows that the overall activity of the variants is in agreement with the log enrichment measured in the bacterial assays (Figure 2B, Figure S6), indicating that the bacterial high-throughput assays quantitatively reflect intrinsic deamination activity.

### Editing at endogenous loci in HEK293T cells is often limited by Cas9-dependent factors or locus accessibility and not by ABE intrinsic activity

We then wanted to test the editing of the ten selected variants at endogenous loci in human cells. They were fused to either nSpCas9 or nSpRY-Cas9, and were tested for activity at several validated editing sites in HEK293T cells (CTNNB1, hs267, HBG1, VEGFA3)^2^ (Figure 2C, Figure S7A). At the peak of the editing window (A4-A5), we observed an increase of editing as the variants’ activity increases. Editing then plateaus and any additional activity does not lead to higher editing. However, the editing of bases outside of the most edited positions keeps increasing after the editing at the peak is saturated (Figure 2C, Figure S8). In addition, the saturation threshold was different for different loci and for different Cas9 orthologs, with PAM-broadened nSpRY-Cas9-fused variants plateauing at lower editing levels across all sites. Although different target sites may reach different plateaus, as activity increases the general trend of saturation is the same. These data suggests that the maximum editing efficiency is often not limited by the intrinsic deaminase activity, but by other factors that could include target search and R-loop formation by the Cas protein and chromatin accessibility of the sites.

The overall trend can be summarized by the mean editing at each window position within the spacer across the tested loci (Figure 2D). This trend shows an increase of the window width as the activity of the variants increases. Wide editing windows are not desired, because they lead to bystander mutations and compromise the single nucleotide precision of an ABE. In addition to A bystanders, we also observed low, but detectable levels of cytidine (C) deamination within the editing window, also increasing with the variants’ activities (Figure S7B).

The observed saturation behavior is consistent across the tested loci suggesting that after a certain threshold, any additional activity does not contribute to increase of the targetable edit but leads to bystander editing and a widening editing window.

### High deaminase activity leads to toxicity in high-throughput assays

In some bacterial assays, we noticed that variants with high activity unexpectedly had low enrichment values. This led us to hypothesize that high deaminase activity could induce mutational burden and genotoxicity, confounding assay results. We therefore introduced another variable in our assays: duration of expression. If the base editors are induced for a long time, we reasoned that their overall enrichment would depend not only on the reversion of antibiotic resistance, but also on the toxicity due to the accumulation of mutations in the bacterial genome or transcriptome (Figure 3A). We assayed our designed library in the two types of selections– easy and hard–and with short versus long induction time, and observed that the highly active variants show a quantitative depletion (Figure 3B). To confirm that the toxicity was not due to the PAM-less nature of SpRY-Csa9, we tested our library with SpCas9 and observed the same drop in enrichment at long induction (Figure S9), supporting the hypothesis that we are measuring properties solely dependent on the deaminase domains of the ABEs.

**Figure 3.**
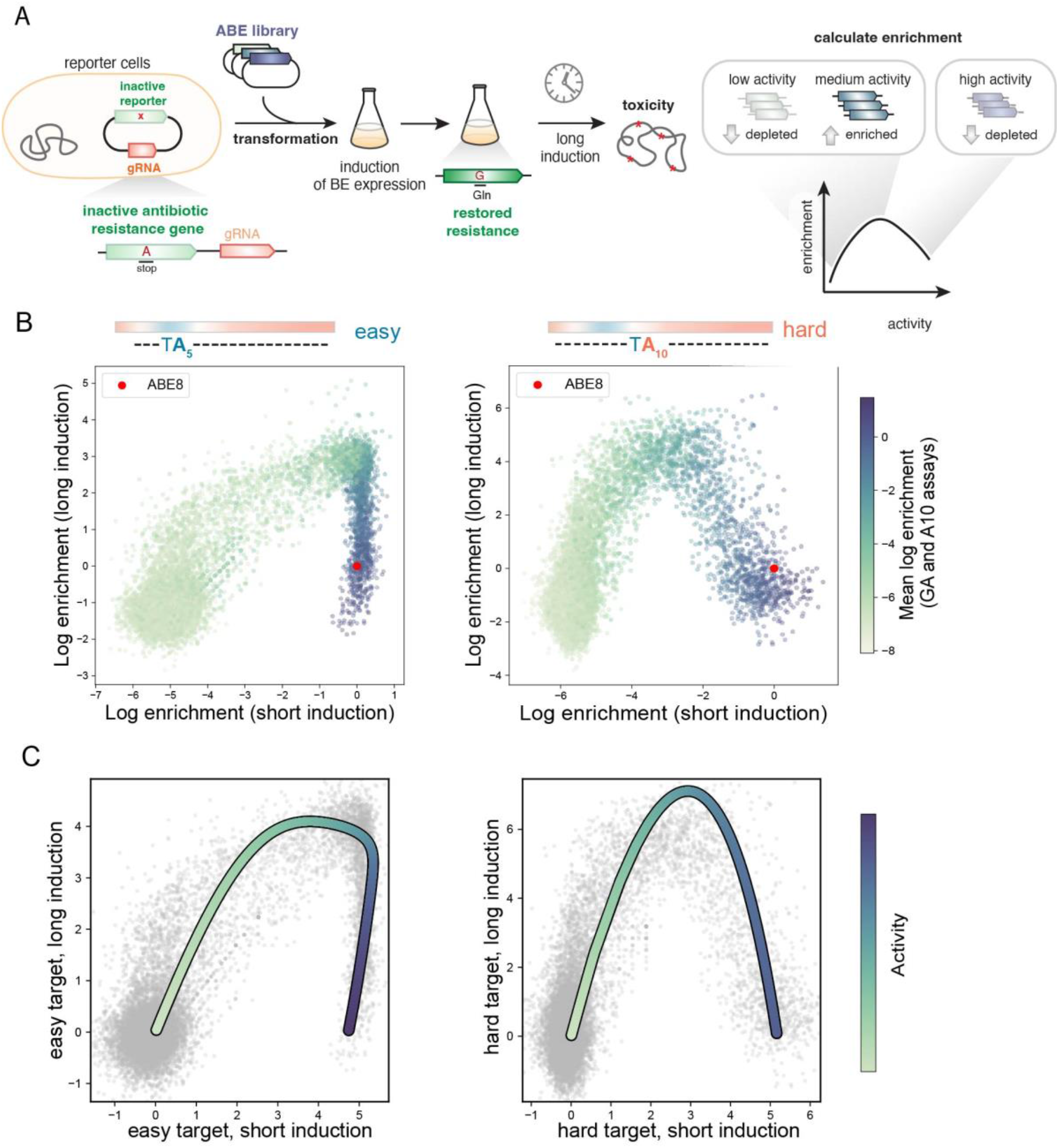
Toxicity in bacterial selections is related to ABE activity. A. Proposed mechanism of bacterial selections with long induction times of ABE expression. Continuous expression leads to toxicity due to spurious DNA mutations across the genome, that can be characterized by the particular shape of their enrichment patterns, as variants with high activity would show lower enrichment. B. Correlations of activity at short and long induction times for two different types of assays (easy and hard). C. Example model enrichment curves. The model can produce a wide variety of pairwise enrichment curves that are parametric with respect to the intrinsic deaminase activity. The curves shown demonstrate some of the complex features seen in the library as exemplified by the data in B. Across all assay conditions the library variants cluster along these apparent one dimensional curves in which the progress along the curve is determined by activity alone.

To explain this behavior, we developed a simple model wherein the on-target activity is used to track the ensemble probability of cured plasmids determining cell survival, and the off-target activity produces non-specific genome edits that accumulate and exponentially decrease the growth rate (see Methods). Crucially, there is only one intrinsic activity, and the on-target and off-target editing probabilities are directly proportional to this single dimension. Systematically varying the proportionality constant for the on-target activity (reflecting the difficulty of the target) and the induction time, we can explore the space of enrichment curves between any pair of simulated assay conditions (Figure S10). Each such curve is parametric in the intrinsic activity, with its characteristic shape governed by the interplay of survival benefit and genotoxic cost. The shape of these curves for the two assay conditions (easy and hard) at long induction times match the shapes of the experimentally measured log enrichments (Figure 3C).

More broadly, the model qualitatively recapitulates the full variety of curve shapes seen across our nine assay conditions using a single axis of activity (Figure S10). These assays span distinct targets, differing in the preceding nucleotide and in editing window position (and hence in intrinsic difficulty), as well as a range of induction times that set the cells’ cumulative exposure to genotoxicity. To test this hypothesis of one-dimensionality independently of the model, we fit a principal curve directly through the nine-dimensional enrichment data (Figure S11). A single one-dimensional curve through this space explains 92% of the variance in the data — essentially all of the reproducible signal, set by the replicate noise ceiling at 93% — and far more than the 82% captured by the best straight line (Figure S11A). The data are therefore confined, within experimental noise, to a one-dimensional manifold (Figure S11B). Moreover, this coordinate is predictive: a variant’s position along the curve predicts a its enrichment in assays withheld from the fit at R^2^ = 0.89, and only about three assays, picked at random, among the nine are needed to place a variant to this accuracy (Figure S11C,D). The remaining assays add no new information, as expected if they are redundant readouts of one coordinate rather than independent dimensions. Our model identifies this single coordinate as the intrinsic deaminase activity.

### Genome-wide off-target effects increase with ABE activity

We next investigated how different types of off-targets relate to the single dimension of activity. First, we focused on the genome-wide DNA off-targets. Previously, an orthogonal R-loop based assay has been shown to reflect the propensity of ABEs to edit non-cognate sites by opening an R-loop with an orthogonal Cas9 *in trans* ^*28*^. We leveraged this idea to build a high-throughput selection on the same principle as our on-target assay, but instead of the ABE being targeted to the antibiotic resistance gene by its fused SpRY-Cas9, an orthogonal dSaCas9 is recruited to the rescue site and the ABE-SpRY-Cas9 library is paired with a non-targeting guide. The resulting rescue directly measures the off-target editing at a non-cognate site (Figure 4A). We pre-selected the library for on-target activity to deplete non-functional variants prior to challenging it with this orthogonal R-loop selection. Up to a certain threshold, we didn’t observe any enrichment in the off-target assay, but as the activities increase, the measured off-target R-loop editing correlates positively with the on-target activity (Figure 4B). The orthogonal R-loop assay and the long induction condition selections are independent methods to capture the off-target editing: in the R-loop assay the off-targets show as higher enrichment (Figure 4A), and in the long induction activity assay the off-targets are measured by the loss of enrichment (Figure 3B). When the R-loop assay data is compared to the long induction time points (for both easy and hard selections) we observe a negative correlation above the detection threshold for the R-loop assay (Figure S12A). Importantly, both of these high-throughput assays show a strong correlation between off-target editing and activity. Additionally, these two methods allowed us to inspect outliers from one assay across the other assays, confirming that there are no systematic outliers in the off-target assays (Figure 4D, Figure S12B), consistent with our observations of one-dimensionality of the ABE activity.

**Figure 4.**
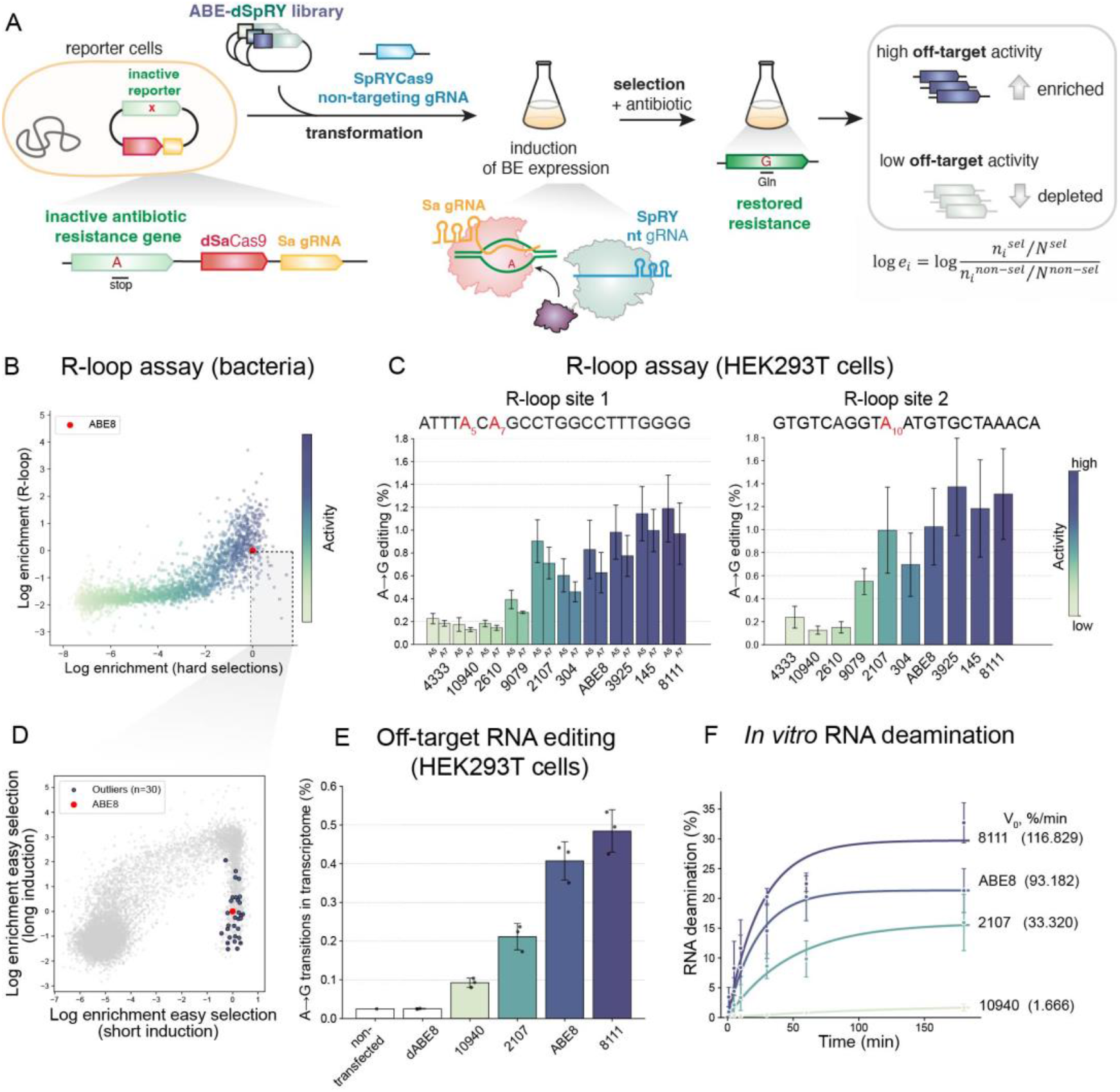
Global DNA and RNA off-targets of ABE variants. A. Schematic of the R-loop selection assay. The inactivating mutation in the antibiotic resistance gene is targeted by a SaCas9 guide, and the ABE library is fused to dSpRY-Cas9 which has a non-targeting guide (nt). The rescue under antibiotic challenge is a readout of the off-target activity of the ABE variants. B. Bacterial R-loop assay selection of the ML-designed library plotted against the mean log enrichment from the two hard selections. The library tested in the R-loop assay was preselected to contain only active variants. A positive correlation between the on-target and R-loop off-target activity is seen with the high activity variants. Variants are colored by their mean log enrichment of the two hard assays. C. Orthogonal R-loop assay of selected variants in HEK293T cells. Bars represent mean editing of three biological replicates, each averaged from three technical replicates. Error bars show standard deviation of the biological replicates. D. Variants that appear to have high activity and low R-loop activity in the R-loop assay are highlighted on one of the long induction assays (easy selection, A5), showing that they have high toxicity. E. Off-target transcriptome-wide A-to-I editing in cellular RNA in HEK293T cells measured by RNAseq. F. *In vitro* RNA deamination rates measured at single-turnover conditions with purified deaminase variants. RNA deamination is measured with EndoV-coupled assay. Quantification of three separate reactions is shown, with errors showing standard deviation for the three independent replicates.

To validate this in HEK293T cells, we tested our selected variants in an orthogonal R-loop assay that measures the off-target editing in mammalian cells concurrently with on-target editing^28^. We observed the same trend of increasing off-target editing with the variants’ on-target activity (Figure 4C). Meanwhile, the on-target editing measured at the targeted site in the HEK293T genome (site 2) follows the same pattern as shown earlier (Figure 2C), with effective editing reaching a saturation (Figure S12C, D). These effects create an illusion that specificity is improved near the saturation point and decreases beyond it. However, the apparent specificity gain is fully attributable to reduced intrinsic deaminase activity and does not reflect a fundamental underlying change in editing selectivity.

### Off-target RNA editing is also dependent on deaminase activity

ABEs’ predecessor is an RNA-editing enzyme TadA, and efforts have been made to minimize RNA editing while improving DNA editing capabilities. However, transcriptome-wide editing has been observed for early engineered variants ^13,15,29^ and ABE8 has been reported to have relatively high RNA editing^2^. In order to characterize the RNA editing of our variants we took parallel *in vivo* and *in vitro* approaches. We selected a subset of variants that spanned the full activity range, transfected them in HEK293T cells and measured the transcriptome-wide RNA edits (Figure 4E, Figure S13). There was a similar level of RNA editing by ABE8 as previously shown^2^, and the variants that we chose showed a strong trend of increasing RNA off-target editing as their activity increases. We also tested them in an *in vitro* EndoV-coupled assay using a ssRNA substrate (Figure 4F, Figure S14). The RNA editing was overall lower than the corresponding DNA editing, suggesting that RNA is a worse substrate, but the RNA editing increased with the deaminase activity. Overall, we conclude that RNA off-target editing is also directly dependent on the ABEs’ deaminase activity.

### Cas9-dependent off-target editing increases with the deaminase activity

Another type of off-targets can also be dependent on Cas9, due to its tolerance to mismatches between the guide and the spacer. Since ABEs require only R-loop formation by the Cas protein to expose the ssDNA, they lack the additional allosteric regulation tied to the conformational changes leading to Cas9 cleavage and are more permissive to edits at mismatched spacers ^30,31^. We tested the activity of our variants at two sites (site 3 and 5) with three known off-target sites for each of them^2^ (Figure 5A). We observed editing at these sites which followed the same pattern as the on-target editing. Additionally, they had the same bystander editing efficiency in cases where there was more than one editable A within the window (Figure S15A). We did not observe a strong trend of the on-target to off-target ratio for the variants, as this ratio seemed to be different for the different sites (Figure S15B,C). When averaged across the off-target sites, there is a small increase of the on-target to off-target ratio at the low activity variants (Figure S15D,E). The overall trend of increasing off-target activity, as the on-target activity increases, was also true for these Cas9-dependent off-target sites.

**Figure 5.**
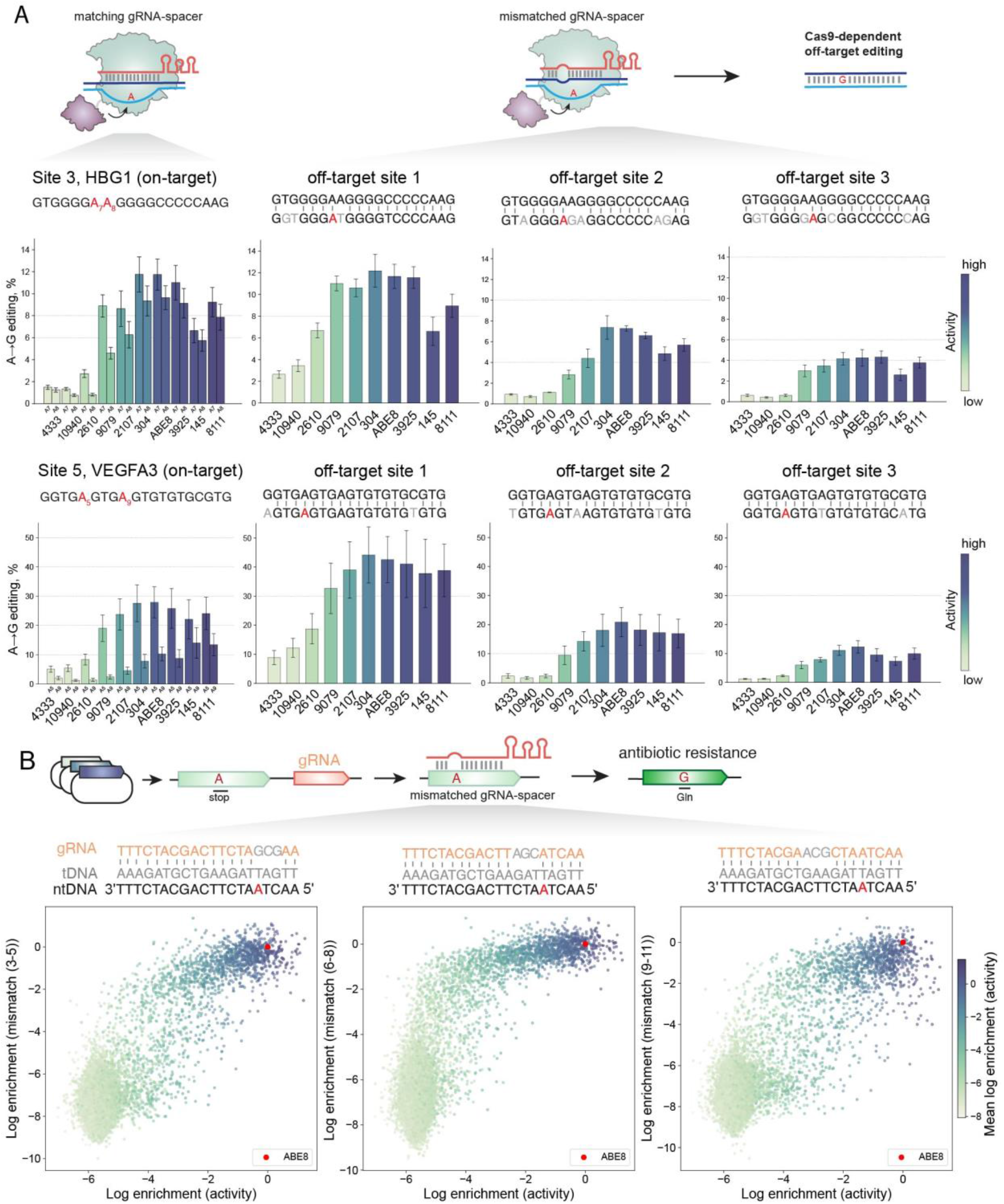
Cas9-dependent off-targets of ABE variants. A. Editing at two HEK293T loci (HBG1, site 3, and VEGFA3, site 5): left, on-target editing of selected variants, right: editing at three known off-target sites is shown for each of the on-target sites. Variants are colored by their activity measured by mean log enrichment in the two hard bacterial assays. Bars represent mean editing of three biological replicates, each averaged from three technical replicates. Error bars show standard deviation of the biological replicates. B. High-throughput selections of the ML-designed library in three different selections that contain tiled three mismatches between the guide and the target DNA. The target is a spacer containing a stop codon in the beta lactamase gene on the reporter plasmid (easy selection 2). Plotted are the correlations between their activity in the hard assay (A10) and the three mismatched selections, showing saturation of the rescue across all mismatched guide targets.

We wanted to understand what the general tolerance for mismatched R-loops of our library variants was and whether it would also be dependent on the activity. We designed three different bacterial reporter strains, containing three tiled mismatches in the guides targeting the antibiotic resistance gene (Figure 5B). Assay of the library in these off-target assays showed that high-activity variants had high levels of editing in all three assays. We were expecting that these targets would be more difficult to edit and would correlate with the hard selection conditions, but they showed a saturation behavior, meaning that the active variants can easily tolerate mismatches. This confirms that depending on their activity, ABEs can edit mismatched targets in a bacterial context with similar efficiency as cognate sites.

### Variants with higher specificity have lower on-target activity

So far, we have observed that in our designed variants there are no exceptions to the rule that increasing specificity is only possible by decreasing the overall activity. A limitation of our approach is that the sequence diversity is focused on a particular region of ABE and it is possible that mutations outside of this region or mutations on residues that we kept fixed in our library can break this dependency. To test this, we assayed additional variants that have previously been reported to have improved specificity and contain mutations both inside and outside our redesigned region, including some of the positions we did not diversify (Figure 6A). We chose a subset of literature variants that have been reported to have either narrower editing windows, lower RNA off-target or lower DNA off-targets^13,15,18,19,23,24^. We introduced all the mutations on top of an ABE8 background (with the exception of HK which is built on an ABE7 background, as per design^24^). When we tested the on-target activity in the bona fide activity assay, all literature variants had reduced activity compared to ABE8 (Figure 6A, Figure S16). In our off-target assays, long induction and orthogonal R-loop assay, we saw that the majority showed low off-target effects and low toxicity (Figure 6C,D), but all were positioned on the one-dimensional curve governed by their activity. With the exception of K20A_R21A, they did not show activity in the orthogonal R-loop assay, nor had a drop of activity at the long induction time point, which is likely due to their low overall activity (Figure 6C).

**Figure 6.**
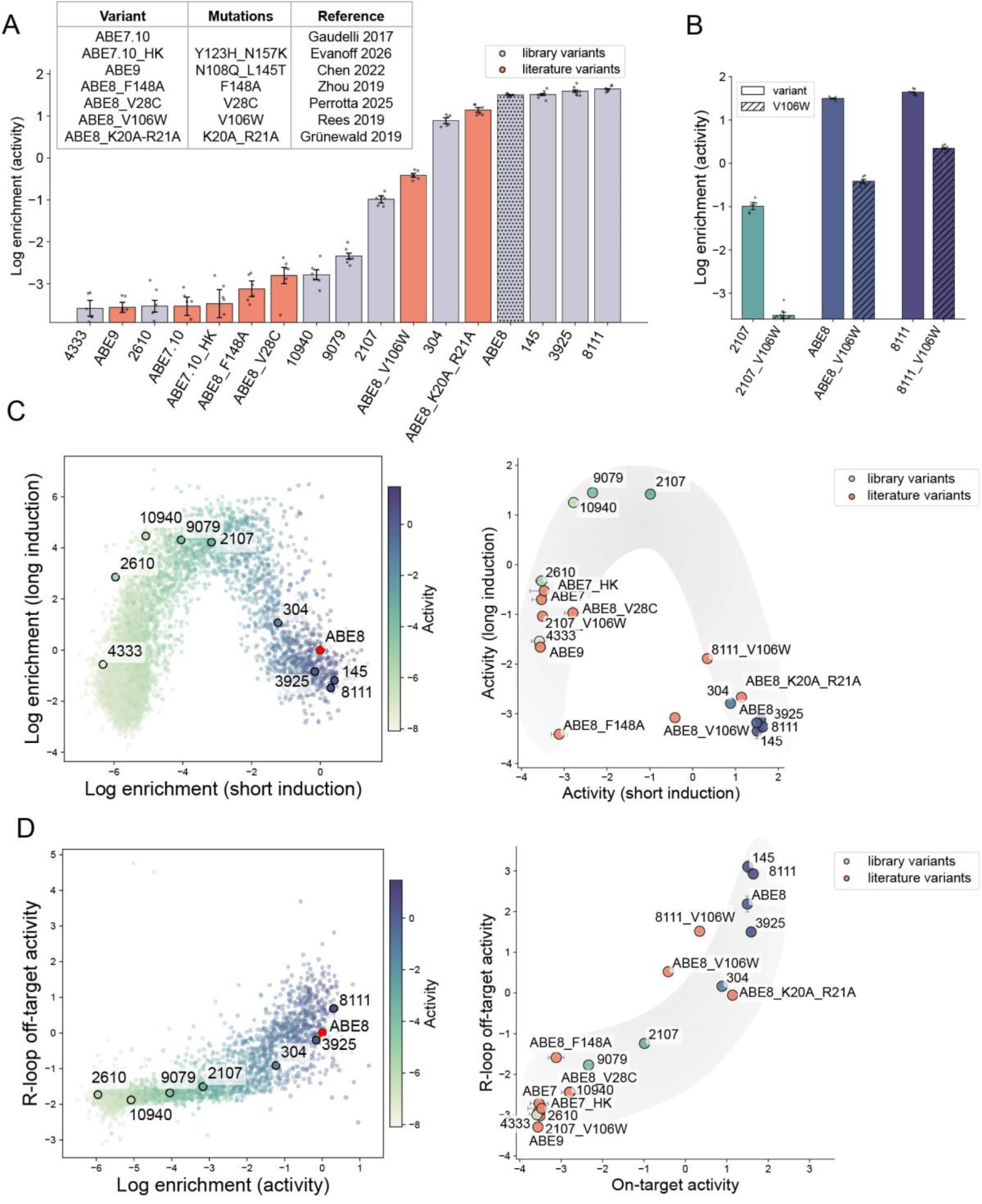
Characterization of previously reported variants with higher specificity. A. Activity measurements from on-target selection of selected library (gray) and literature-reported (orange) variants. The variants were tested as a pool in one of the hard selections (A10) with a 2h induction time. B. Effect of the V106W mutation on ABE8, 2107 and 8111 variants from the same pooled assay, compared to the corresponding variants without the mutation. C. Correlation between short and long induction as a measure of the off-target induced toxicity. Left: position of the selected variants in the full library assays. Right: log enrichments from the pooled experiment of library and literature variants at short (2h) and long (16h) induction times with the A10 selection. D. Variants’ log enrichments at the activity assay (as measured by the 2h induction time in A10) and the R-loop assay. Left: position of the selected library variants in the library experiment, right: pool of library and literature variants in the activity and R-loop assays.

We also considered V106W, which has been used in multiple studies including a recent clinical study^9,15,32–35^, as an important mutation, and introduced it on three different backgrounds—ABE8 and two variants from our library with lower and higher than ABE8 activity (2107 and 8111). V106W decreased the activity of all three variants it was introduced on - ABE8, 2107 and 8111 (Figure 6B), strongly suggesting that the mechanism of action of this mutation, and potentially all the other mutations, is to decrease the activity under the saturation threshold for mammalian target editing, where there is no obvious loss of on-target editing, but a noticeable decrease in off-target editing.

## Discussion

Our data shows that the primary functional dimension of the deaminase domains of ABEs is their intrinsic activity, and both on-target and off-target editing are linked to this property. An important observation we made is that increasing the deaminase activity leads to increase of the editing efficiency at the optimum of the editing window at its target site, up to a ceiling, after which no increase of the editing is effectively observed (Figure 7A). When this plateau is reached, higher activity manifests as more off-target editing.

**Figure 7.**
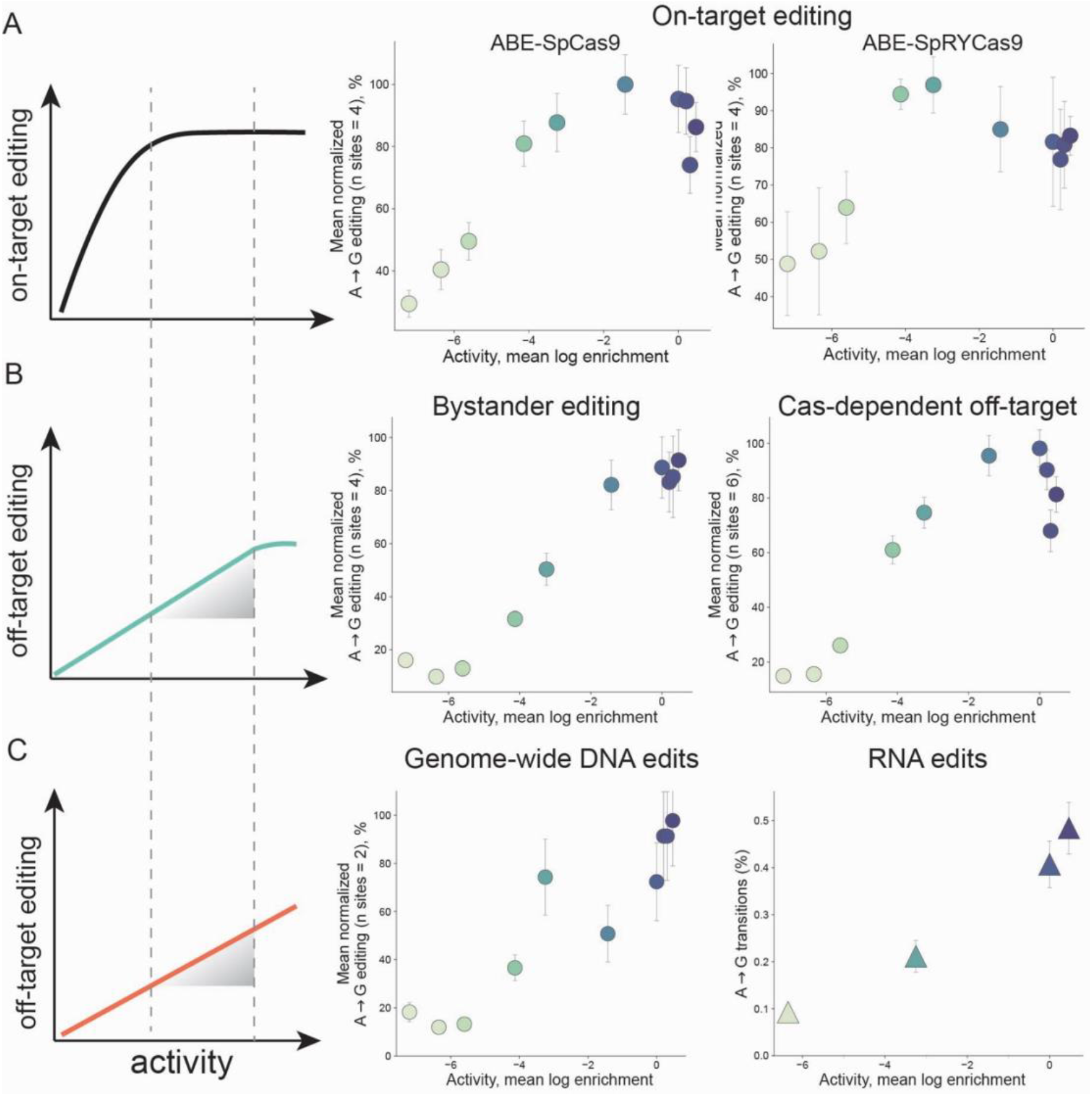
Trends of on-target and off-target editing by ABEs. A. On-target editing has a saturating pattern, increasing intrinsic deaminase activity leads to increase of the effective editing, but after a threshold additional activity does not translate to higher editing. The limit is both site- and Cas9-dependent, but the saturation shape is universal. Left: schematic of the editing saturation, middle and right: mean normalized editing at peak positions for 4 sites for SpCas9 (middle) and SpRY-Cas9-fused ABE variants (right). Peak window positions are as follows, site1: A4, site 2: A5, site 3: A7, site 5: A5. For all plots the activity on the x-axis is shown as measured by the mean log enrichment of the two hard high-throughput assays shown on Figure 1D. B. Some types of off-targets (bystander edits and Cas9-dependent off-target edits) also have a saturating behavior, but the rates at which the limit is reached are different than the peak-window on-target editing. Left: schematic; middle: mean editing at a chosen bystander position for 4 sites for ABE-SpCas9 variants (site 1: A9, site 2: A8, site 3: A8, site 5: A9); right: mean normalized editing at peak positions of 6 off-target sites (site 3, off-site1 and 2: A7, off-site 3: A8; site 5, off-site1, 2 and 3: A5). C. Genome and transcriptome-wide edits are theoretically not limited and keep increasing as the deaminase activity increases. The ratio of on-target to off-target editing can therefore look different for variants with different activities, but it is a function of the intrinsic deaminase activity. Left: schematic; middle: mean normalized orthogonal R-loop activity at the two measured sites (site 1: A5, site 2: A10); right: RNA off-target editing measured by RNAseq. All sites are normalized to their own maximum editing so they could be averaged. Error bars represent ±1 standard deviation across three biological replicates, propagated across sites as the square root of the summed per-site variances divided by the number of sites, after normalizing each site to its maximum.

A classical perspective of enzyme specificity defines it as the apparent catalytic efficiency for a substrate over the catalytic efficiency for a similar, but structurally distinct molecule^36^. In the case of ABEs, specificity is viewed in terms of the combined specificity of the Cas9 and the deaminase domain. From this perspective, the ABE specificity is the editing at a desired position within the editing window, over other, “bystander” positions (within the same window) or over other DNA loci, not targeted by Cas9.

At the R-loop formed by Cas9, the substrate of the deaminase is any adenine within the single - stranded non-target DNA. This means that additional bystander A’s are equivalent substrates for the deaminase and they would also get edited. In the case of the parent TadA enzyme, the substrate specificity is likely determined by the precise structure of its tRNA anticodon loop substrate^37,38^. The accommodation of an extended DNA substrate during the ABE evolution has been accompanied by an opening of the substrate-binding site that has decreased the structural specificity of the interaction with the nucleic acid^27^. While this is a prerequisite for enabling the programmability of the ABEs, it has likely compromised their specificity, e.g. the interaction with the substrate must be sequence-independent and any structural constraints would limit it. An exception of that is the observation that all ABEs have an inherent preference for a pyrimidine at the −1 position preceding the target A^1^. Increasing the tolerance for a preceding G coincides with an increase of overall activity (Figure 1E). In the case of perfectly programmable ABE, the preference of some positions likely arises from structural constraints imposed by the position of the deaminase fused to Cas9. Therefore, the editing at secondary positions would also increase together with the deaminase activity, and theoretically would also reach saturation, but at a different rate (Figure 7B). Effectively, this means that after the editing at the peak has reached a ceiling, any surplus activity will just serve to widen the editing window (Figure 2D).

Outside of the Cas9-targeted R-loop, any exposed ssDNA in the genome is also a potential substrate for ABEs. This gives rise to two types of off-target edits: Cas9-dependent and Cas9-independent edits. The first type are DNA off-targets that are caused by transient opening of non-cognate R-loops by Cas9, where there is a partial match between the gRNA and the target DNA. These Cas9-dependent off-targets often possess the highest editing rates, and for some sites they can be as high as the on-targets. The fact that the ABEs can edit those sites as highly as they edit their fully cognate sites, suggests that the deamination is not limiting in this scenario, and modulating the properties of the ABE domain is unlikely to improve this type of specificity, as their editing would also increase with the deaminase activity (Figure 7B).

Another example of DNA off-targets are genome-wide edits. It is not well understood whether these edits are caused by transient binding and formation of short-lived R-loops during the dCas9 search, or whether ssDNA substrate is exposed during cellular processes such as replication or transcription. However, any ssDNA that is accessible could be edited by the deaminase. While we have measured this type of Cas9-independent off-targets with an R-loop assay, which is a proxy, we propose that increasing the activity of the ABEs would directly lead to higher off-target edits throughout the genome (Figure 7C).

One notable exception of ABEs’ substrate specificity is the preference of DNA over RNA. However, the opening of the active site to accommodate the extended DNA conformation has enabled editing of ssRNA^27^, and we and others have observed that an increase in DNA editing activity is accompanied by an increase of RNA editing (Figure 7C)^2^. We propose that, from a theoretical framework, it is possible to increase the specificity of DNA over RNA, potentially by imposing structural specificity for the deoxyribose over the ribose rings.

We show that there is an inherent trade-off between activity and specificity and even highly diverse ABEs do not escape it. Moreover, previous attempts have not engineered ABEs out of this strict activity and specificity relationship. It is therefore likely that the current class of TadA-derived ABEs are fundamentally limited in their precision. From a practical perspective we propose navigating this trade-off by choosing an ABE variant with just sufficient activity to approach the editing ceiling, since any excess activity will serve only to increase off-target edits. Another factor for regulating the exposure of the targeted DNA is the PAM recognition by Cas9. PAM binding is important for both positioning the ABE at a precise location in the window, and for the R-loop formation, therefore affecting the accessibility of the DNA substrate^39,40^. We also observed that variants with different PAM preferences affect the overall editing by the ABEs. Using a combination of Cas9 variants with different PAM requirements and deaminase with a minimal activity required to reach the editing threshold can help increase the ratio of desired versus non-desired edits ^41^. With our ML-based design method we were able to produce numerous deaminase variants spanning a wide range of activities. In practice, one could select from among these for an activity best matched to the delivery modality, induction, etc. in order to achieve adequate editing while minimizing, to the extent possible, damaging excess. Fundamentally altering the activity and specificity trade-off, however, will likely require architectural innovations to push the next generation of ABEs to greater functional precision.

## Methods

### General cloning

All plasmids were assembled using NEB HiFi or Golden Gate assembly. PCRs were performed with Q5 (NEB) or PrimeStar (Takara) polymerases, fragments were DpnI (NEB)-digested and gel-extracted or PCR cleaned-up with DNA Clean and Concentration Kit (Zymo). NEBuilder HiFi assembly (NEB) was performed at 50°C for 1h; for Golden Gate, fragments were incubated at 37°C for 1h with corresponding Type IIS enzyme (BsaI-HF, NEB) and T4 DNA ligase (NEB). For general cloning the assembled DNA was transformed in Turbo cloning cells (NEB). Plasmids were prepared with QIAprep Spin Miniprep Kit (Qiagen) and sequences were verified with full-plasmid nanopore sequencing.

### Machine learning-guided library design

#### Probabilistic Sequence Model Used

We scored candidate sequences with one overall Potts model whose energy (unnormalized log-likelihood) was the average of two different Potts models—one learned from evolutionary information in the form of a Multiple Sequence Alignment (MSA), and the other from structural information (backbone-sequence pairs) by distilling an “inverse-folding” model (a model that provides a distribution of sequences conditioned on a backbone structure) into a Potts model. As Potts models, each has a probability distribution over sequences that is a function of an energy, *E*(*x*), such that *p*(*x*) ∝ exp(−*E*(*x*)), where *x* = (*x*_1_, …, *x*_L_) is a sequence of length *L* = 167 (the TadA deaminase domain) a 21-letter alphabet (20 amino acids plus gap),

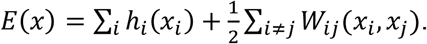

The lower the energy, the more favorable (probable) the sequence. The two models differ only in how their parameters were estimated (i.e., the fields *h* and couplings *W*), described next. The advantage of this combined model over either single modality model alone was validated on deep-mutational-scanning data (Figure S1A) and in prior work^42^.

#### Evolutionary model

The evolutionary model was trained on an MSA of natural TadA homologs generated following the EVcouplings protocol^43^. Starting from wild type *E. coli* TadA (167 residues) as the query, we ran two iterations of jackhmmer^44^ against UniRef100^45^ with a bitscore inclusion threshold of 0.5 bits per residue (83.5 bits), applied to both full sequences and individual domains. The resulting alignment was converted to a2m format, retaining match-state (uppercase) columns and mapping insertions to gaps, yielding 73,037 homologs over 167 match columns. Sequences with 30 or more gaps were discarded, and each remaining sequence was weighted by the inverse of the number of sequences within 80% identity to it (computed over aligned, non-gap positions). The ABE8 deaminase sequence was added to the weighted alignment with a weight of 1 prior to fitting.

Model parameters were estimated by maximizing the sequence-weighted pseudolikelihood of the alignment using stochastic gradient descent (learning rate 0.1, 100 epochs, minibatch size 1000, sequences sampled in proportion to their weights), with an *L*_2_penalty:

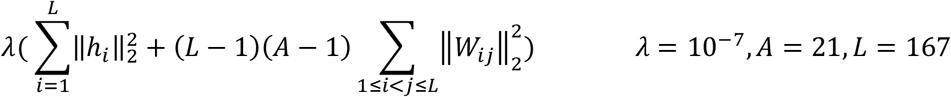

The (*L*−1)(*A*−1) factor scales the coupling penalty relative to the fields and was originally proposed by^42^. We denote the learned energy of this Potts model as *E*_evo_(*x*).

#### Inverse-folding model distilled into a Potts model

We started with the inverse-folding model ESM-IF1 (esm_if1_gvp4_t16_142M_UR50^46^), an autoregressive model which allows us to compute the negative log-likelihood (NLL) of any sequence conditioned on a fixed backbone (here, the backbone used was the AlphaFold-Multimer prediction for the ABE8 deaminase homodimer). Because computing the ESM-IF1 negative log-likelihood is too slow to query inside our sampling-based design procedure described, we first distilled it into a Potts model whose energy can be evaluated cheaply for large batches of sequences. As our design objective (below) depends only on the unnormalized Potts log-likelihood, we do not need the normalizing constant, making these models fast and cheap for our use case.

We explored two distillation strategies. In the first, we sampled sequences from ESM-IF1 conditioned on the ABE8 backbone and fit a Potts model to these samples by maximum pseudolikelihood, capturing the model’s global sequence distribution. In the second, we set the Potts parameters directly to exactly reproduce the ESM-IF1 NLLs of: the wild type sequence, all single mutants, and all double mutants. Specifically, choosing the wild type gauge, we can set the “field” and “coupling” parameters as follows, denoting *wt* as wild type, *wt*_{*i*→*a*}_ as the wild type with mutation to amino acid *a* at position *i*, and similarly, *wt*_*{i→a,j→b}*_denoting a double mutation

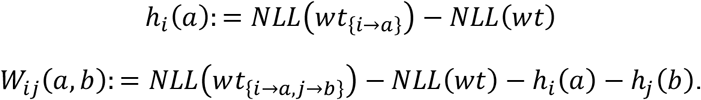

It may be helpful for some readers to view this Potts model as a second-order expansion of the ESM-IF1 score around the wild type, thereby yielding a model whose fit is most accurate near the wild type sequence. This strategy requires computing the ESM-IF1 NLL for every single double mutation. Thus, for computational efficiency in the second strategy, we employed the approximation that any two residues not in structural contact (inter-atomic distance ≥6.0 Å, including the dimer interface) had coupling parameter values equal to zero, and hence could be omitted from the computation; this approximation sharply reduces the number of ESM-IF1 evaluations required. Both distillation strategies reproduced ESM-IF1 scores on held-out sequences well; we chose to use the second (local-fit) model for library design because its distillation should be more accurate near the wild type. We denote the learned energy of this Potts model as *E*_struc_(*x*).

#### Overall Potts Model

We combined the two aforementioned Potts models into a single Potts model by averaging the two model’s energies,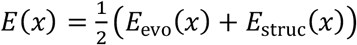. Because the Potts energy is linear in its parameters, we can equivalently average their parameters, namely the fields and couplings,

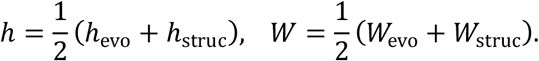

This overall energy, *E*(*x*), was used to score all sequences for all library design (Figure 1A, Figure S1B).

### Library Design

#### Overview and design groups

Using the overall Potts energy as a fitness surrogate, we designed a library of ABE8 deaminase domain variants by sampling sequences that were high-scoring (low energy). However, because the highest-scoring library has no diversity, we explored the trade-off between higher average fitness and higher diversity, generating a new library for each possible trade-off (Figure S1B)^47^. Diversity was defined as the average pairwise hamming distance of every two sequences in the library. The designed region comprised residues 33–118, containing the active site and the central part of the protein; all positions outside this window were held fixed at the ABE8 wild type sequence. However, we generated three design groups that differ in which positions within this window were allowed to vary, and the maximum number of mutations away from wild type allowed in any library sequence (“mutation cap”):

1. **core** — only buried positions were designable (mutation cap of 20 from ABE8);
2. **surface** — only solvent-exposed positions were designable (mutation cap of 20);
3. **core+surface** — both shells were designable (mutation cap of 40).

Buried and exposed positions were defined from the per-residue solvent-accessible surface area (SASA) of the ABE8 AlphaFold model (chain A), computed with biotite^48^; residues with SASA < 5 Å^2^ were classed as core (49 residues) and the remainder as surface (118 residues). Within the design window we additionally froze the catalytically/structurally critical positions identified in prior ABE evolution (residues 48, 74, 106, 108, 109, 111) and the zinc-coordinating residues (57, 59, 87, 90) (Table S1). Mutations to cysteines were not allowed.

#### Objective function

Rather than design sequences independently, we treated the entire batch (the library) as the object of optimization, so that diversity of the library could be directly incorporated into the design^37^. For a library of *B* sequences {*x*^1^, …, *x*^*B*^}, each of length *L*, we defined the mean pairwise Hamming distance as:

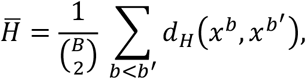

where 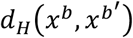 is the hamming distance between sequences *x*^*b*^ and 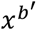. We then minimized the library-level objective:

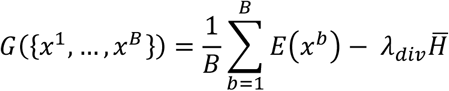

The first term favors sequences with low ensemble energy, while the second favors libraries with high average pairwise sequence diversity. Because the objective is minimized, the negative sign causes libraries with larger mean pairwise Hamming distance to be preferred. To select the diversity penalty weight, λ_*div*_, for all three libraries we traced out the energy-diversity trade-off curve by scanning increasing values of λ_*div*_ (Figure S1B) and heuristically chose a single value that appeared to balance low-energies with a high enough diversity for all three design groups, 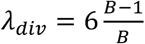.

We note that 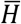 can be computed efficiently without explicitly summing over all pairs of sequences. For a pair of sequences *x*^*b*^and 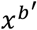, the Hamming distance can be written as:

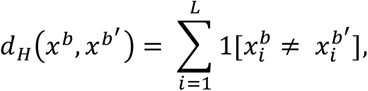

where 1[·]denotes the indicator function. At each position, the mismatch indicator is the complement of the agreement indicator:

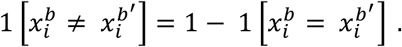

Therefore,

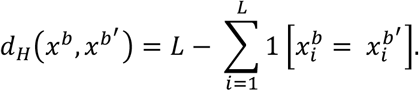

Averaging over all pairs gives:

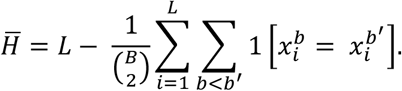

Let 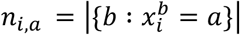 denote the number of sequences containing amino acid *a* at position *i*.

Because 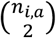is the number of sequence pairs that both contain amino acid *a* at position *i*,

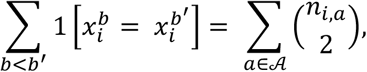

where *A* is the set of 20 standard amino acids. Substituting the above expression gives:

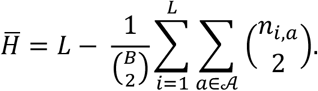

This expression allowed us to evaluate the mean pairwise Hamming distance from per-position amino-acid counts rather than by explicitly comparing every pair of sequences.

#### Parallel simulated annealing

We minimized the objective over the whole library using a batched, gradient-informed Markov-chain Monte Carlo sampler^49^ under simulated annealing. All *B* sequences were initialized at the ABE8 wild type. At each step, the gradient of the temperature-scaled joint objective, *G*({*x*^1^, …, *x*^*B*^}), with respect to the one-hot sequences was used to form, for each sequence, a categorical proposal distribution over single-residue substitutions, with each candidate scored by its first-order predicted change in the objective. A substitution was proposed and then accepted or rejected by a Metropolis–Hastings test (including the forward/reverse proposal correction) for every sequence in the batch in parallel.

We note that this parallel update is an approximation. Because the diversity penalty couples all sequences in the batch through the shared per-position counts *n*_i_,_a_, the objective is a single function of the entire library, and a strictly correct Metropolis–Hastings chain would update one sequence at a time — evaluating each proposal against the fixed current state of the rest of the batch. Updating all sequences simultaneously does not exactly preserve detailed balance for this joint distribution. Due to the large size of the library we are designing (*B* = 5,000), we adopt this approach for computational efficiency and rely on the sequences being only weakly coupled (each contributes to the penalty only through counts that change by one out of *B* members per substitution), so that the parallel scheme approximates the correct single-sequence updates well and becomes exact in the limit of independent sequences.

Constraints (frozen positions, no new cysteines, and—once a sequence reached its mutation cap—only reversions toward ABE8 wild type) were enforced within the proposal by masking disallowed substitutions with −∞ logits, so they were never proposed. Annealing was implemented by dividing the objective by a temperature *T* initialized at 0.9 and multiplied by 0.99 at each of 200 steps (final *T* ≈ 0.12), shifting the sampler from broad exploration toward low-energy, high-diversity configurations. After 200 steps, the batch was deduplicated to unique sequences and the 4,000 lowest-energy unique designs were retained (see above).

Our goal was to have B=4,000 unique sequences in one design group. Because the optimization of the objective function can yield libraries with duplicate sequences, for each design group we actually designed 5,000 sequences, and then deduplicated these 5,000 sequences, by keeping the 4,000 lowest-energy (best) unique. Keeping 4,000 for each of the three (core, surface, core+surface) yielded a total of 12,000 designs spanning 1–40 mutations from ABE8.

#### Library cloning

A Golden Gate destination vector for the library was designed with a pSC101 origin of replication and the ABE-Cas9 placed under an arabinose promoter. Designed libraries containing overhangs for PCR amplification with a total length of 300 bp were ordered from Twist biosciences. The 12,000 sequences were divided in three sub-libraries based on their number of mutations from ABE8 (1-10, 11-20, 21-40) ordered with separate overhangs and cloned separately. PCR of the libraries was performed with KAPA HiFi polymerase (Roche) using 2 ng template and 200 nM primers for 20 cycles with 1 min extension time to reduce cross-over products. The product was run on a 2% agarose gel and gel-extracted with Zymoclean Gel DNA recovery kit (Zymo) and used for Golden Gate assembly (NEB) into a destination vector containing a GFP sequence flanked by BsaI sites at the region where the designed library is inserted. The Golden Gate assembly was incubated at 37°C for 5 min and 16°C for 5 min, cycled through 40 times, with a final incubation at 37°C for 1h and heat inactivation at 65°C for 5 min. Reactions were cleaned up with DNA Clean and Concentration Kit (Zymo) and transformed in Top10 cloning strain (NEB) via electroporation with a BioRad Micropulser. After electroporation, cells were supplemented with 950 µL SOC media and incubated at 37 ºC for 1h in a shaking incubator at 200 rpm. 10-fold dilutions were made of the transformants and were plated to ensure the transformation efficiency exceeds at least 10 times the number of expected variants in each sub-library. The rest of the outgrowth after the recovery was used to inoculate 50 mL 2xYT media supplemented with 35 µg/mL chloramphenicol, cultures were grown for 12-14h and DNA was purified with QIAprep Spin Miniprep Kit (Qiagen). The sublibraries were mixed in a ratio based on the number of variants in each sublibrary to have a final theoretically equimolar amount of each variant.

### Bacterial high-throughput selections

#### Reporters and selection strains

Bacterial selections were built based on a p15A origin reporter plasmid containing a constitutively expressed inactivated carbenicillin resistance gene and a guide RNA targeting the inactivating mutations (Supplementary sequences, Table S2). Top10 cells were transformed with the reporter plasmid and competent cells carrying the plasmid were prepared. Briefly, reporter plasmid were transformed in Top10, a single colony was used to inoculate 1 L culture using 2xYT media and plasmid maintenance antibiotic (kanamycin). The culture was grown to OD600 of 0.6 at 37°C in a shaking incubator. Cells and reagents were kept on ice for the full procedure. The cultures were cooled down and centrifuged for 10 min at 4000 g at 4°C. All the supernatant was removed and pellets were washed by resuspending them in sterile cold water. The water wash step was repeated twice, followed by a wash in 10% glycerol. After the glycerol wash cells were centrifuged at 4000 g for 13 min at 4°C and resuspended in 2.5 mL of 10% glycerol (for 1 L starting culture), aliquoted in 50 µL, flash-frozen in liquid nitrogen and stored at −80°C.

#### Selections

The plasmid library was transformed in the reporter strains by adding 200 ng plasmid in 100 µL competent cells. Cells were incubated on ice for 5 min and electroporated using a BioRad Micropulser and immediately supplemented with 900 µL SOC media. The cells were incubated at 37 °C on a shaking incubator at 200 rpm for 1h. After the recovery 4 mL 2xYT media with maintenance antibiotics (chloramphenicol and kanamycin) was inoculated with the transformed cells (1 mL) and induced with 0.1% arabinose for the indicated time (Table S3). At the end of the induction the cells were centrifuged at 4000 g for 10 min, resuspended in 1 mL 2xYT media and centrifuged again for 10 min to wash away any inducer. The resulting pellet was resuspended in 1 mL 2xYT and inoculated in 29 mL 2xYT with 400 µg/mL carbenicillin and 35 µg/mL chloramphenicol (selective conditions) and 50 µg/mL kanamycin and 35 µg/mL chloramphenicol (non-selective conditions). Cultures were grown incubated at 37 °C at 200 rpm until reaching OD600 of 0.6. For the long induction conditions, 30 µL of the 1 mL resuspended pellet were inoculated in the 30 mL media with antibiotics. After reaching OD600 of 0.6 cells were centrifuged at 4000 g for 10 min and plasmid DNA was isolated with Qiagen miniprep kit.

100 ng/µL of the isolated DNA post selection was used for PCR with primers carrying Illumina adaptors (PCR1). Samples were sent for PCR2 and sequencing.

For the experiments with selected subset of variants and literature variants, all plasmids were cloned separately and an unique barcode was added on the plasmid, with three different barcodes per variant. The selections were performed identically, but instead of the BE region, the unique barcode was amplified and prepared with native-ligation protocol for nanopore sequencing.

#### Analysis of bacterial selections

Each bacterial selection assayed a pooled library of base-editor variants under a selective condition (antibiotic challenge) and a matched non-selective condition in independent biological replicates. Variant abundances were measured by amplicon sequencing of the variable region before and after selection, and activity was quantified as the enrichment of each variant under selection relative to the non-selective condition. The same pipeline was applied to all bacterial selections in this work; individual assays differ only in the selective condition imposed (e.g. the required edit position or sequence context, the orthogonal R-loop construct, guide mismatches, or the induction time).

#### Read processing and variant counting

For each sample, paired-end reads were merged into a single consensus sequence spanning the variable region. The constant sequences flanking the variable region were located in each mate by a minimum-mismatch search and used to orient and register the read pair; the overlapping portion of the two mates was reconciled base-by-base by taking the higher base-quality (Phred) call, and the merged read was trimmed to the fixed-length window corresponding to the designed region. Each merged sequence was matched exactly to the set of designed library members; reads without an exact match to any library member were discarded. This yielded, for every sample, a table of read counts per designed variant.

#### Enrichment quantification

For each replicate, the per-variant count tables from the selective and non-selective conditions were combined, and a pseudocount of one was added to all counts to stabilize ratios for low-count variants. Counts in each condition were converted to relative frequencies (a variant’s count divided by the total counts in that sample), and the log enrichment of each variant was computed as

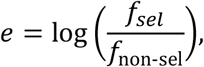

the log-ratio of its frequency under the selective versus non-selective condition. Log enrichments were then referenced to ABE8 by subtracting the ABE8 variant’s log enrichment within each replicate, so that *e* = 0 corresponds to ABE8 activity.

#### Deaminase protein purification

ABE variants were cloned in an expression vector containing a His_14_-SUMO tag on the N-terminus, followed by an ABE monomer. Expression plasmids were transformed in BL21-DE3 Lobstr strain (Kerafast), single colonies were inoculated and grown overnight at 37°C in a shaking incubator at 200 rpm. Saturated cultures were used to inoculate large scale expression (1 to 6 L). Cells were grown to OD600 of 0.6 and induced with 1 mM Isopropyl β-D-1-thiogalactopyranoside (IPTG), followed by expression at 16°C for 16 h. Cells were harvested by centrifugation at 4000 g for 30 minutes and pellets were flash frozen in liquid nitrogen. The cells were resuspended in lysis buffer containing 50 mM HEPES NaOH pH 7.5, 500 mM NaCl, 10 % glycerol v/v, 1 mM Tris(2-carboxyethyl)phosphine (TCEP), C0mplete protease inhibitors (Roche, 1 tablet per 50 mL lysis buffer). The lysate was sonicated for 5 min total time at 70 % amplitude with 5 sec on and 15 sec off pulse on a Qsonica sonicator. Lysate was clarified by centrifugation in a JA20 rotor for 30 min at 30,000 g and the soluble fraction (supernatant) was loaded on 1 mL NiNTA resin, equilibrated with buffer A (50 mM HEPES NaOH pH 7.5, 500 mM NaCl, 10 % glycerol v/v, 20 mM imidazole, 1 mM TCEP). The resin was washed with 10 column volumes of buffer A, followed by 10 column volumes of buffer B (buffer A with 1 M NaCl), 10 column volumes of buffer A and eluted with 1 column volume fractions of buffer E (50 mM HEPES NaOH pH 7.5, 500 mM NaCl, 10 % glycerol v/v, 400 mM imidazole, 1 mM TCEP) until all protein was eluted (followed with Bradford reagent and/or A280 absorption on a nanodrop). Elution fractions were loaded on a 4-20 % gradient SDS-PAGE and fractions containing protein of interest were pooled. The SUMO tag was cleaved with His-tagged SUMO protease in dialysis overnight in buffer containing 50 mM HEPES NaOH pH 7.5, 250 mM NaCl, 10 % glycerol v/v and 1 mM TCEP. After cleavage, the protein was loaded onto 1 mL Ni-NTA resin and the flow-through was collected and centrifuged for 10 min at 20,000 g in a JA20 rotor. The protein was then diluted in buffer C (50 mM HEPES NaOH pH 7.5, 10% glycerol v/v and 1 mM TCEP) to a final NaCl concentration of 30 mM and loaded on HiTrap SP cation exchange column (Cytvia) on an AKTA Pure FPLC system. After loading, the column was washed with one column volume buffer C supplemented with 30 mM NaCl and eluted with a 30 mL gradient from 30 mM to 1 M NaCl (3 % to 100 % buffer D, 50 mM HEPES NaOH pH 7.5, 1 M NaCl, 10 % glycerol v/v and 1 mM TCEP). The eluted peak was concentrated with Amicon Ultra centrifugal filters (10 kDa cutoff), followed by centrifugation for 5 min at 21 000 g and the sample was loaded on Superdex S75 10/300 GL (Cytiva) size exclusion column in 50 mM HEPES NaOH pH 7.5, 150 mM NaCl, 5 % glycerol v/v and 1 mM TCEP. Peak fractions were pooled and concentrated with Amicon Ultra centrifugal filters (10 kDa cut-off) to a final concentration of 50-150 µM, flash frozen in liquid nitrogen and stored at −80°C. Protein concentrations were measured by A280 absorbance using an extinction coefficient of 18571 M^−1^ cm^−1^ (calculated by ProtParam).

#### *In vitro* DNA editing assay with direct inosine detection

To date all *in vitro* measurements of TadA deaminase kinetics have been indirect, mostly via gel-based characterization of cleavage products after treatment by an inosine-specific endonuclease^2,26,27^. Below we describe a new method that couples biochemical assay with nanopore sequencing of synthetic DNA substrates to directly detect inosine from its effect on the electrical current during strand translocation (Figure S5). We trained a custom model that identifies adenosine versus inosine at a defined site using the raw nanopore signal. Many reaction conditions including different variants, kinetic timepoints, and replicates can be designated with defined barcode ligation, pooled for nanopore amplicon sequencing, and demultiplexed post-facto to determine adenine editing efficiencies in high throughput.

#### DNA preparation

DNA editing by the deaminase domains was measured on a 60-nucleotide single stranded DNA substrate (ssDNA) (Supplementary sequences). Substrate and complementary PAGE-purified DNA were purchased from IDT. Complementary DNA was purchased with 5’ phosphorylation for the ligation reaction. Barcodes were purchased as separate oligonucleotides. The 5’ of the forward strand was phosphorylated with T4 PNK (NEB) in T4 ligase buffer for 1h at 37 ºC, followed by quenching at 65 ºC for 20 min. The phosphorylated barcodes were purified with Oligo Clean and Concentrator Kit (Zymo) according to the manufacturer’s protocol. Concentrations were measured on a nanodrop at A260. Extinction coefficients were calculated based on their sequence with the IDT OligoAnalyzer− Tool Oligo Calculator. The phosphorylated oligos were annealed prior to ligation with equimolar amounts of their complementary strand by heating up to 95 ºC and cooling down at a rate of 0.1 ºC/s to 20 ºC.

#### DNA deamination reactions

For the deamination reactions, 5 µM enzyme was incubated with 1 µM ssDNA in reaction buffer (50 mM Tris-HCl pH7.5, 5 mM NaCl, 5 mM MgCl_2_, 10 µM ZnCl_2_, 5% glycerol v/v, 1 mM TCEP). The deaminases were stored in SEC buffer (50 mM HEPES NaOH pH 7.5, 150 mM NaCl, 5% glycerol v/v and 1 mM TCEP) and were all diluted to 50 µM stocks in the SEC buffer, so the same volume could be added to each reaction. Reactions were initiated by adding substrate and incubated at 37 ºC for 1, 5, 10, 30, 60 and 180 min. To quench the reactions, 10 µL of the reaction mixtures were mixed with 5 µL complementary DNA (56 nucleotides) at a final concentration of 0.67 µM to have an equimolar ratio and incubated at 95 ºC for 10 min. The complementary DNA contained two mismatches opposite of the edit position, so it could be distinguished at the sequencing step. The resulting double stranded DNA with a staggered end was ligated to the pre-phosphorylated barcoded dsDNA. For the ligation, 0.67 µM dsDNA barcode with a complementary staggered end were attached to each time point sample using 400 units T4 DNA ligase (NEB) in T4 ligase buffer for 16 h at 16 ºC, followed by heat inactivation at 65 ºC for 10 min. Samples from different time points, each containing a different barcode, were pooled and purified on a DNA Clean & Concentrator 5 column (Zymo). The pools of barcoded substrates were then prepared for sequencing using the Oxford Nanopore native ligation procedure that incorporates additional sample barcodes and sequencing adapters for recognition by the nanopore helicases and sent to the UC Berkeley Barker Sequencing facility.

#### Model training for inosine basecalling

The default nanopore basecaller Dorado (Oxford Nanopore) does not recognize inosine so we had to train a custom model. Defined synthetic substrates of length 60 bp were ordered with either an adenosine or inosine at the edit site and hybridized with DNA that was fully complementary except for the edit and adjacent upstream positions. These double-stranded synthetic substrates were submitted for native ligation nanopore amplicon sequencing.

Dorado was used for unmodified base calling of the reads using the “emit moves” flag mapping the base calls to the raw signal. Bowtie2 was used to align the reads to the reference substrate sequence using the “very-sensitive-local” flag since the actual reads extend beyond the substrate sequence with barcode and adapter sequence. From these alignments the forward reads, i.e. those containing the edit base, were selected along with each corresponding raw current signal pod5 entry. The modified base calling software package Remora (Oxford Nanopore) was trained on this ground-truth data with adenosine and inosine while manually withholding 10% of the reads for subsequent model validation. The training data contained 43,384 control (A) reads and 27,250 inosine reads. We used the LSTM model architecture with 8 layers (--size 8) and a chunk context size of 100 (--chunk-context 100) for training. We noticed that the basecaller transition time calls at the inosine had higher variability than at canonical nucleotides and reasoned that binary classification on the raw signal directly might enhance the accuracy. To this end, a lightweight custom convolution neural net (CNN) was trained on the raw signal centered on the edit position with +/−100 points. On the same training data the CNN achieved similar performance to Remora. However, combining the models by averaging their read call probabilities we were able to substantially improve the performance with the ensemble model. See the table for model performance metrics showing the false positive (FP) and false negative (FN) rates for the different models used.

| Model | FP (raw) | FN (raw) |
| --- | --- | --- |
| Remora ta_lstm_s8_c100 | 4.5% | 3.4% |
| custom CNN | 4.0% | 3.0% |
| ensemble | 2.9% | 2.7% |

#### Read demultiplexing and analysis

The raw pod5 files for the multiplexed substrate reads (see “DNA deaminase reactions”) were basecalled by the default Dorado model while emitting moves with alignment to the unedited substrate sequence. After filtering for forward reads, Remora inference and the CNN trained above were used in ensemble to predict the probability of adenosine to inosine editing. Reads with >50% predicted chance of inosine were counted as edited. Given a table mapping the ligated barcodes to each reaction condition, the reads were demultiplexed using cutadapt, and the results were aggregated. For each condition the reported edited fraction was computed as,

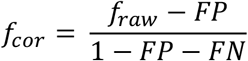

With *f*_*cor*_ as the corrected edited fraction, *f*_*raw*_ as the observed raw edited fraction, *FP* as the model false positive rate, *FN* as the model false negative rate. Any values below 0 or exceeding 1 were clipped to 0 and 1, respectively. Measurements were repeated in triplicate with the points shown as the mean and whiskers for +/−one standard deviation.

The resulting product concentration versus time data for each variant were fit to a single-exponential association model:

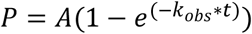

where A is the fitted plateau amplitude and *k*_*obs*_ is the observed first-order rate constant, using nonlinear least-squares regression. Initial reaction velocities (v0) were calculated from the fitted parameters as:

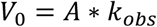

corresponding to the initial slope of the exponential model at *t* =0.

#### *In vitro* RNA editing assay

RNA editing was measured with an Endonuclease V-coupled assay^27^. The RNA substrate was purchased from IDT, EndoV enzyme was purchased from NEB. The substrate sequence was chosen with the criteria to contain only a single A and to have no predicted secondary structures at the reaction temperatures. For the deamination reaction, 5 µM purified deaminase variants were incubated with a 30-mer 5’-FAM-labelled RNA substrate (Supplementary sequences). Reaction buffer contained 50 mM Tris-HCl pH7.5, 5 mM NaCl, 5 mM MgCl_2_, 10 µM ZnCl_2_, 5% glycerol v/v, 1 mM TCEP. All enzymes are diluted from 50 µM stocks in SEC buffer (see DNA deamination reactions). The enzyme and reaction buffer were pre-mixed and the substrate RNA added to a final concentration of 1 µM to initiate the reaction. Reaction mixtures were incubated at 37 ºC for 1, 5, 10, 30, 60 and 180 min. At each time point, 10 µL of the reaction mixtures were quenched at 95 ºC for 10 min. 5 units of EndoV diluted in NEBuffer4 were added to the quenched reaction mixture and incubated at 37 ºC for 1h, followed by heat inactivation at 65 ºC for 20 min. The resulting reactions were incubated at 95 ºC for 3 min in loading buffer containing: 95% formamide, 0.025% SDS and 100 mM ethylenediaminetetraacetic acid (EDTA). The samples were run side by side with a buffer sample containing the same loading buffer supplemented with 0.025% bromophenol blue to follow the sample migration during electrophoresis. The samples were loaded on a 15% denaturing TBE-Urea gel, run at 100 V for 90 min in TBE buffer. The gels were imaged on a GelDoc XR+ and bands were quantified with FIJI. Product formation was calculated as the fraction of cleaved DNA and fitted to a single exponential model, identical as for the DNA editing (see *In vitro* DNA editing assay with direct inosine detection).

### HEK293T cell transfections and editing at endogenous loci

#### Transfections

HEK293T cell transfections were performed following previously established procedure^2^. Cells were maintained by passaging at lower than 80% confluency and passaged up to passage 22. They were kept in DMEM, high glucose, GlutaMAX Supplement, pyruvate media (ThermoFisher Scientific) supplemented with 10% FBS (Gibco, lot number 2565838RP) and trypsinized with TrypLE. Cells were negative for mycoplasma by ATCC mycoplasma test. Plasmids for transfections were purified with QIAGEN Plasmid Plus Midi Kit.

Cells were seeded at 50,000 cells per well in a 48-well Clear Flat Bottom TC-treated Cell Culture Plates (ThermoFisher Scientific) 24h prior to transfection. Transfections were performed with Lipofectamine 3000 (ThermoFischer Scientific). 750 ng ABE-nSpCas9 or ABE-nSpRYCas9 plasmid and 250 ng guide plasmid (total 2 µL) were premixed with 2 µL P3000 reagent with 12.5 µL Opti-MEM™ I Reduced Serum Medium (ThermoFischer Scientific). 1.5 µL Lipofectamine 3000 reagent were mixed with 12.5 µL Opti-MEM medium and added to the DNA-P3000 mixture. The resulting mixture was incubated for 20 min at room temperature and 30 µL were added per well. Cells were incubated with the DNA-Lipofectamine mix for 48 hours.

For all transfections, master mixes of 3x were prepared and three wells were transfected for technical replicates. The triplicate transfections were performed three times on separate days at different cell passage for biological replicates.

The orthogonal R-loop experiments were performed identically, except the DNA mixture contained 375 ng of dSaCas9 plasmid, 250 ng SaCas9 guide plasmid, 375 ng ABE-nSpCas9 plasmid and 250 ng SpCas9 guide. The cells were incubated with the DNA-Lipofectamine mix for 72 hours.

After 48 h (or 72 h for the orthogonal R-loop assay, respectively) media was removed and cells were lysed with 100 µL lysis buffer (10 mM Tris-HCl pH7.5, 0.05% SDS, 25 µg/mL Proteinase K). The 48-well plates were incubated with the lysis buffer at 37 ºC for 15 min to detach the cells and were then transferred to a PCR plate, followed by incubation for another 45 min (for a total of 60 min) at 37 ºC and heat inactivation of the Proteinase K at 80 ºC for 20 min. Lysates were diluted 1:10 in DEPC-treated H_2_O and used to prepare samples for NGS sequencing. Samples were barcoded with the PCR primers, so they could be cleaned up in a pooled fashion. Primers for amplification of the different target sites contained barcodes for demultiplexing and Illumina adaptors. PCR1 reactions contained 0.4 µM forward and reverse primers, 1 µL diluted lysate and Q5 2x MasterMix (NEB) in 25 µL reactions. Samples with different barcodes were pooled and cleaned up with DNA Clean and Concentrator Kit 5 (Zymo) and sent for PCR2.

#### Demultiplexing and analysis of HEK293T editing

The pooled and barcoded paired end reads of the HEK293T editing site amplicons were demultiplexed using cutadapt (v5.0) ^50^ with trimming in order to identify the variant, condition, and replicate for all reads and to remove the barcodes for further analysis. CRISPResso2 was used in batch mode to quantify the editing efficiencies across each of the demultiplexed samples ^51,52^. The following parameters were used: --base_editor_output TRUE, -s (minimum single base quality score) 10, -wc (editing quantification window center position) −10, -w (editing quantification window half width) 10, --conversion_nuc_from A, --conversion_nuc_to G. The CRISPResso editing results were consolidated and the means and standard deviations for the replicates were computed position-by-position for each tested variant, site and condition combination.

### Transcriptome-wide RNA editing in HEK293T cells

#### Transfections for RNA editing

For RNAseq experiments cells were seeded in 24-well plates at 100,000 cells per well 24 h prior to transfection. 1500 ng ABE-nSpCas9 and 500 ng guide plasmids were mixed with 4 µL P3000 reagent and 25 µL Opti-MEM medium. 3 µL Lipofectamine 3000 were mixed with 25 µL Opti-MEM and added to the DNA-P3000 mixture, followed by incubation at room temperature for 25 min. 60 µL of the transfection mix were added to each well and incubated for 48 hours. Media was removed and RNA was purified with RNeasy kit (Qiagen) following the manufacturer’s protocol. 350 µL RLT buffer was added to the wells to resuspend the cells and transferred to a Qiashredder column, followed by spinning down at 21,000 g for 2 min. The flow-through was collected and transferred to a new tube. 300 µL 70 % ethanol was added and carefully mixed by pipetting. The mix was transferred to a RNeasy column, centrifuged at 10,000 g for 30 sec, the flow-through was discarded. 350 µL RW1 buffer was added, centrifuged at 10,000 g for 30 sec and flow-through discarded again. 10 µL DNase I was mixed with 70 µL buffer RDD, gently mixed and applied to the column and incubated for 15 min at room temperature. 350 µL of RW1 buffer were added, centrifuged at 10 000 g for 30 sec and flow-through discarded. The column was washed twice with 500 µL RPE buffer, centrifuged at 10 000 g for 30 sec for the first wash and 2 min for the second wash. The column was transferred to a new collection tube, centrifuged for 2 min at 21,000 g and eluted with 30 µL RNase-free H_2_O for 1 min at 10,000 g. Concentrations were measured on a nanodrop, normalized to 80 ng/µL and sent for RNAseq with Plasmidsaurus.

#### RNA-seq analysis

The data consists of single end reads with the original RNA sense enriched for mRNA sequences and biased to the 3’ due to poly-A specific amplification and further deduplicated by UMI to correct for PCR artifacts. The aligned bam files were filtered to only the main chromosomes and indexed with samtools.

REDItools3 (v3.6)^53,54^ was used to identify and count RNA mutations using the following options: --strand 1 (identifying the correct sense of the RNA with respect to the reads), --min_edits 0 (recording all positions, even those with no observed mutations), --min_read_depth 5 (only recording positions with a depth of at least 5 reads). The REDItools3 strand inference doesn’t work properly with single end reads so we applied a small patch. To limit the effect of false positives due to SNPs, sites having greater than 20% mutation and 5 or more reads for any of the negative controls (i.e. the catalytically dead ABE8 or the non-transfected sample) were blacklisted and excluded from downstream quantitative analysis. The adenine deamination fraction was computed at all non-blacklisted A’s as G /(G+A).

#### Modeling on-target activity and off-target toxicity of base editors

Motivated by the ABE library assay results we constructed an analytical model to better understand the interplay between on-target editing, which enables survival through antibiotic resistance repair, and off-target editing, which leads to accumulating toxicity via diffuse genome edits. We use this model to explore the null hypothesis that any variant is described by just a single intrinsic deamination activity through which the on- and off-target activities are rigidly linked.

#### On-target activity

The analytical model of on-target editing activity simulates the probabilistic events of target base editing of eligible unedited plasmids and the replication and random partitioning of plasmids into daughter cells during cell division. In our treatment we make the assumptions that the plasmid copy number is uniform, the replicated plasmids are divided equally between daughters, and there is no repair of DNA mismatches at the editing site—that is, the base editor only makes edits to one strand to make a “half-edited” plasmid and subsequent resolution of the mismatch occurs only through replication which produces one unedited and one fully edited copy.

To track the evolution of the probability distributions of edited plasmids, we use a stochastic matrix formalism. Given an initial normalized probability vector, **p**_**0**_, multiplication by the stochastic matrix, **P**, yields the updated probability vector of plasmid edit numbers, **p**_**f**_, through a single cell doubling cycle. The terms of **P** can be exactly expressed using a hypergeometric distribution for plasmid division weighted by a binomial distribution of editing outcomes across all eligible plasmids.

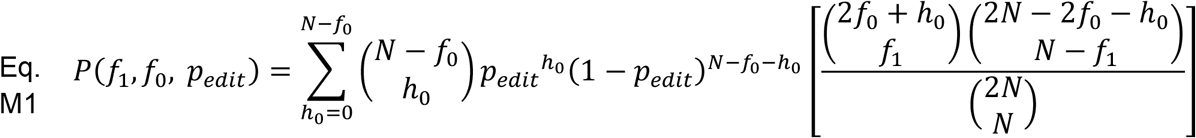

Here *f*_*0*_ is the initial number of edited plasmids, *f*_*1*_ is the number of plasmids edited after the update, *p*_*edit*_ is the probability that a single unedited plasmid is edited during the cell doubling time, *N* is the plasmid copy number, and h_0_ is the number of half-edited plasmids that serves as a dummy index of summation. The terms of form 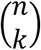 represent combinatorial factors, *n* choose *k*, and use the convention that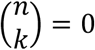 for *k > n*.

Each element of the stochastic matrix is the probability that *f*_*1*_ plasmids are edited given an initial state of *f*_*0*_ edited plasmids for one cell doubling. For repeated doublings the stochastic matrix is raised to the power of the number of doublings and multiplied through initial probability vector. We verified that the analytical formula in Eq. M1 matched numerically simulated base editing outcomes.

Finally, to go from the plasmid edit probability distribution to a cell-by-cell survival probability we need a survival vector, **s**, the elements of which represent the probability that a cell with a given number of edited plasmids can survive the antibiotic selection. For example, if a single edited plasmid is sufficient then **s** would be 0 followed by *N* ones, meaning that it cannot survive without any edits but can with 100% probability with one or more edited plasmids. The overall probability of survival is the weighted sum of the survival vector weighted by the final edit probability vector which is simply the dot product. Thus, the complete expression for the average survival probability as a function of *p*_*edit*_ and the number of doublings in induction, *d*, is,

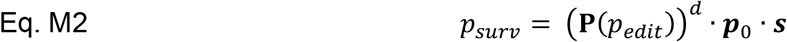

#### Off-target activity

For estimating the relative growth defect arising from off-target genomic edits by the base editor we assume that individual mutations fall into three main types: harmless mutations with no growth consequence, lethal mutations, and mutations having a distribution of small growth defects. The net effect of *k* mutations is then taken as the product of *k* samples from this joint distribution (Fig. M3).

Among the non-lethal mutations we assume that the displacements below one are typically small, i.e. *v*_*i*_ = 1 − ϵ_*i*_ where ϵ_*i*_ is small. Thus, we can approximate the distribution quite generally as an exponential decay with respect to ϵ_*i*_ by recognizing that 1 − ϵ_*i*_ as the first two terms in the Taylor expansion of 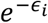. This allows us to convert the product into a summation by the properties of exponentials.

For estimating the relative growth defect arising from off-target genomic edits by the base editor we assume that individual mutations exert, on average, a small decrease in relative fitness (ϵ_*i*_) such that the relative growth rate of a given mutant is *v*_*i*_ = 1 − ϵ_*i*_. Furthermore, we assume that these fitness decreases stack in a multiplicative way as a product over *k* mutations. We can approximate the growth distribution quite generally as an exponential decay with respect to ϵ_*i*_ by recognizing that 1 − ϵ_*i*_ as the first two terms in the Taylor expansion of 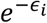. This allows us to convert the product into a summation by the properties of exponentials,

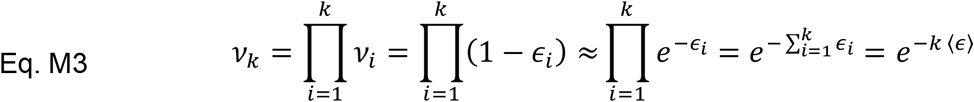

In this formulation we can see that the expected growth rate decreases exponentially with the number of random mutations. We can then attach a specific meaning to the average fitness cost term ⟨ϵ⟩ as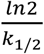 where *k*_*1/2*_ is the number of mutations for which the average growth rate is half that of wild type.

Now we must calculate the average growth rate by performing a weighted average across the number of mutations. We expect that the number of non-specific mutations is Poisson distributed and that expected rate of mutation will scale in direct proportion to the off-target activity constant (*a*_*off*_) and the length of induction (*t*_*ind*_) such that the probability of observing *k* mutations is given by,

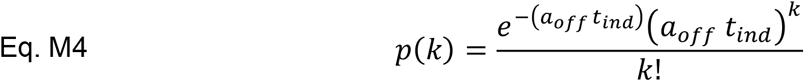

To compute the growth rate we average the rate as a function of *k* in Eq. M3 over the probability distribution of *k* mutations in Eq. M4 which yields a simple closed-form solution.

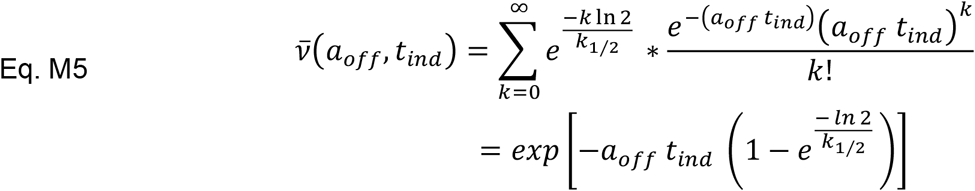

#### Predicting enrichment outcomes

Using our expressions for the survival probability from on-target activity and the growth defect introduced by off-target activity we can calculate predicted enrichments. The enrichment of any given variant depends upon the on-target activity rate constant (*a*_*on*_); the off-target activity rate constant (*a*_*off*_); the induction time (*t*_*ind*_), i.e. the time over which the base editor is acting; and the outgrowth time (*t*_*out*_), i.e. the time over which the cell population is growing subject to antibiotic selection. Before bringing these all together we first must express the survival probability (Eq. M2) in terms of *a*_*on*_ and *t*_*ind*_.

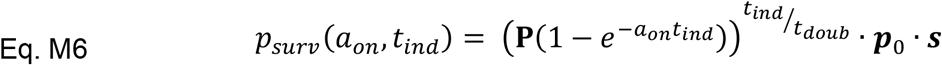

Here, *p*_*edit*_ has been substituted with the time evolution of the editing probability and the doubling number (*d*) is replaced by the induction time divided by the doubling time constant. For non-integer doubling values the matrix power **P**^d^ can be calculated as **P**^*d*^ = **A D**^*d*^**A**^−1^ where **P** has been diagonalized with **A**, the matrix of column-wise eigenvectors of **P**.

Next, we express the expansion of a variant’s population over the outgrowth time (*t*_*out*_) with the assumptions that the viable cells grow exponentially according to their computed average growth rate and that the non-viable cells are static but still contribute their counts (which are with increasing outgrowth time diluted to insignificance by the share of viable cells). *p*_*surv*_ and 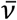are given by Eq. M6 and Eq. M5, respectively.

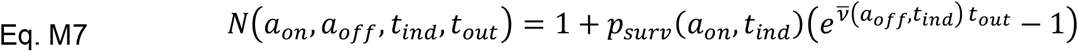

The library selection experiments we seek to model are pooled assays in which the variant enrichments are calculated as the log ratio of the relative counts observed in selection over the non-selective relative counts. Thus, we can write the enrichment of the *i*^th^ variant as,

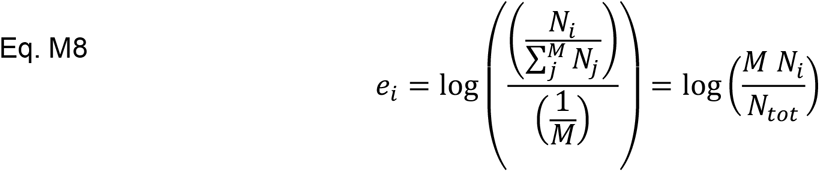

where *M* is the total number of variants and *N*_*i*_ is the outgrowth count of the *i*^th^ variant with *a*_*on*, *I*_ and *a*_*off*, *I*_, *N*_*tot*_ is the sum of all variant outgrowth counts.

The null model condition is that *a*_*on*_ and *a*_*off*_ are not independent and are instead linked via proportionality constants to a common intrinsic deamination activity. Specifically, we assume that each variant has an intrinsic deamination activity, *a*_*i*_, and that this is related to the off-target activity by a single global constant, *a*_*off*,*i*_ = *a*_*i* *_*c*_*off*_. The on-target activity for each variant is also related to the intrinsic activity by a constant, however, since the target sites vary in their editing difficulty according to the preceding base and the position within the editing window, each target will have its own constant, *a*_*on,t,i*_ = *a*_*i*\*_*c*_*on,t*_.

## Author contributions

M.L., L.M.O, H.N., D.F.S and J.L. conceived and designed the study. H.N. designed the library. M.L. did library cloning, bacterial assays, protein purification, *in vitro* assays and mammalian editing experiments. L.M.O. built the on-target and off-target activity model, nanopore sequencing analysis, RNAseq analysis, mammalian cell editing analysis and *in vitro* nanopore-based activity assay analysis. L.M.O., H.N. and M.L. analyzed the data. Y.L. cloned the selected library and literature variants. C.I.T. did preliminary mammalian cell experiments and cloning of mammalian vectors. S.K. assisted with library cloning. C.N. assisted with cloning and preparing competent cells. A.B. assisted with library design. D.F.S. and J.L. supervised the project. M.L. wrote the manuscript with input from all authors.

## Acknowledgments

We thank Brian McCarthy, Scott Geller, Michelle Davila, Julie Vargas and the entire UC Berkeley Barker sequencing facility for nanopore and Sanger sequencing throughout the project. We thank Netravathi Krishnappa and the IGI-NGS core facility for Illumina sequencing. We thank Flora Zhiqi Zhang, Antoine Koehl, Jung-Un Park, Rachel Weissman and the Savage and Listgarten labs for helpful discussions. The research was supported by HHMI. M.L. is a Life Science Research Foundation Fellow, sponsored by HHMI. This material is based upon work supported by the National Science Foundation graduate research fellowship program under grant no. DGE 2146752 (C.I.T.) and 1752814 (A.B.). Any opinions, findings and conclusions or recommendations expressed in this material are those of the authors and do not necessarily reflect the views of the National Science Foundation.

## References

1. Gaudelli, N.M., Komor, A.C., Rees, H.A., Packer, M.S., Badran, A.H., Bryson, D.I., and Liu, D.R. (2017). Programmable base editing of A.T to G.C in genomic DNA without DNA cleavage. Nature 551, 464–471. 10.1038/nature24644.

2. Richter, M.F., Zhao, K.T., Eton, E., Lapinaite, A., Newby, G.A., Thuronyi, B.W., Wilson, C., Koblan, L.W., Zeng, J., Bauer, D.E., et al. (2020). Phage-assisted evolution of an adenine base editor with improved Cas domain compatibility and activity. Nature Biotechnology 2020, 1–9. 10.1038/s41587-020-0453-z.

3. Siegner, S.M., Ugalde, L., Clemens, A., Garcia-Garcia, L., Bueren, J.A., Rio, P., Karasu, M.E., and Corn, J.E. (2022). Adenine base editing efficiently restores the function of Fanconi anemia hematopoietic stem and progenitor cells. Nat. Commun. 13. 10.1038/s41467-022-34479-z.

4. Liao, J., Chen, S., Hsiao, S., Jiang, Y., Yang, Y., Zhang, Y., Wang, X., Lai, Y., Bauer, D.E., and Wu, Y. (2023). Therapeutic adenine base editing of human hematopoietic stem cells. Nature Communications 14. 10.1038/s41467-022-35508-7.

5. Rose, I., Greenwood, M., Biggart, M., Baumlin, N., Tarran, R., Hart, S.L., and Baines, D.L. (2025). Adenine base editing of CFTR using receptor targeted nanoparticles restores function to G542X cystic fibrosis airway epithelial cells. Cellular and Molecular Life Sciences 82. 10.1007/s00018-025-05587-y.

6. Muller, A., Sullivan, J., Schwarzer, W., Wang, M., Park-Windhol, C., Hasler, P.W., Janeschitz-Kriegl, L., Duman, M., Klingler, B., Matsell, J., et al. (2025). High-efficiency base editing in the retina in primates and human tissues. Nat. Med. 31, 490–501. 10.1038/s41591-024-03422-8.

7. Wan, P., Tang, S., Lin, D., Lu, Y., Long, M., Xiao, L., Jiang, Y., Liao, J., Ma, X., Liu, Y., et al. (2026). In vivo base editing gene therapy for heterozygous familial hypercholesterolemia: a phase 1 trial. Nat. Med. 32, 1045–1051. 10.1038/s41591-026-04254-4.

8. Gupta, A.O., Sharma, A., Frangoul, H., Kanter, J., Mapara, M.Y., Dalal, J., Alavi, A., Jaroscak, J.J., Ayala, E., DiPersio, J.F., et al. (2026). Base Editing of HBG1 and HBG2 Promoters for Sickle Cell Disease. New England Journal of Medicine 394, 1824–1835. 10.1056/nejmoa2504835.

9. Musunuru, K., Grandinette, S.A., Wang, X., Hudson, T.R., Briseno, K., Berry, A.M., Hacker, J.L., Hsu, A., Silverstein, R.A., Hille, L.T., et al. (2025). Patient-Specific In Vivo Gene Editing to Treat a Rare Genetic Disease. New England Journal of Medicine 392, 2235–2243. 10.1056/nejmoa2504747.

10. Vafai, S.B., Täubel, J., Ashdown, T., Patel, R.S., Diamondali, S., Cegla, J., Soran, H., Bashir, B., Abitbol, A., Gaudet, D., et al. (2026). In Vivo Base Editing of PCSK9 with VERVE-102 for Hypercholesterolemia. N. Engl. J. Med. 10.1056/NEJMoa2601283.

11. Bower, O.J., R. Orsi, A.E., McMahon, R., Staneva, D., Blagrove, J., Singh, K., Simon, C.S., McCarthy, A., Garcia, P., Shaikly, V., et al. (2026). Base editing reveals an essential role for NANOG in human embryogenesis. Nature. 10.1038/s41586-026-10792-1.

12. Chen, X., McAndrew, M.J., and Lapinaite, A. (2023). Unlocking the secrets of ABEs: the molecular mechanism behind their specificity. Biochem. Soc. Trans. 51, 1635–1646. 10.1042/BST20221508.

13. Zhou, C., Sun, Y., Yan, R., Liu, Y., Zuo, E., Gu, C., Han, L., Wei, Y., Hu, X., Zeng, R., et al. (2019). Off-target RNA mutation induced by DNA base editing and its elimination by mutagenesis. Nature 571, 275–278. 10.1038/s41586-019-1314-0.

14. Grünewald, J., Zhou, R., Garcia, S.P., Iyer, S., Lareau, C.A., Aryee, M.J., and Joung, J.K. (2019). Transcriptome-wide off-target RNA editing induced by CRISPR-guided DNA base editors. Nature 569, 433–437. 10.1038/s41586-019-1161-z.

15. Rees, H.A., Wilson, C., Doman, J.L., and Liu, D.R. (2019). Analysis and minimization of cellular RNA editing by DNA adenine base editors. Sci. Adv. 5.

16. Fiumara, M., Ferrari, S., Omer-Javed, A., Beretta, S., Albano, L., Canarutto, D., Varesi, A., Gaddoni, C., Brombin, C., Cugnata, F., et al. (2023). Genotoxic effects of base and prime editing in human hematopoietic stem cells. Nat. Biotechnol. 10.1038/s41587-023-01915-4.

17. Wu, L., Jiang, S., Shi, M., Yuan, T., Li, Y., Huang, P., Li, Y., Zuo, E., Zhou, C., and Sun, Y. (2024). Adenine base editors induce off-target structure variations in mouse embryos and primary human T cells. Genome Biol. 25. 10.1186/s13059-024-03434-0.

18. Chen, L., Gao, H., Zhong, L., Liu, M., Liu, M., and Cheng, Y. (2022). Engineering a precise adenine base editor with minimal bystander editing. Nat. Chem. Biol. 10.1038/s41589-022-01163-8.

19. Grünewald, J., Zhou, R., Iyer, S., Lareau, C.A., Garcia, S.P., Aryee, M.J., and Joung, J.K. (2019). CRISPR DNA base editors with reduced RNA off-target and self-editing activities. Preprint at Nature Publishing Group, https://doi.org/10.1038/s41587-019-0236-6 10.1038/s41587-019-0236-6.

20. Valdez, I., O’Connor, I., Patel, D., Gierer, K., Harrington, J., Ellis, E., Caponetti, S.A., Sebra, R.P., Valley, H.C., Coote, K., et al. (2025). A streamlined base editor engineering strategy to reduce bystander editing. Nature Communications 16. 10.1038/s41467-025-63609-6.

21. Shang, M., Li, Y., Cao, Q., Ren, J., Zeng, Y., Wang, J., Gonzalez, R.V.L., and Zhang, X. (2025). A motif preferred adenine base editor with minimal bystander and off-targets editing. Nature Communications 16. 10.1038/s41467-025-64203-6.

22. Zhao, N., Zhou, J., Tao, T., Wang, Q., Tang, J., Li, D., Gou, S., Guan, Z., Olajide, J.S., Lin, J., et al. (2024). Evolved cytidine and adenine base editors with high precision and minimized off-target activity by a continuous directed evolution system in mammalian cells. Nat. Commun. 15, 8140. 10.1038/s41467-024-52483-3.

23. Perrotta, R.M., Vinke, S., Ferreira, R., Moret, M., Mahas, A., Chiappino-Pepe, A., Riedmayr, L.M., Mehra, A.-T., Lehmann, L.S., and Church, G.M. (2025). Engineered base editors with reduced bystander editing through directed evolution. Nat. Biotechnol. 10.1038/s41587-025-02937-w.

24. Evanoff, M., Korpal, S., Krill, Z.D., Cowan, Q.T., and Komor, A.C. (2026). Precise, minimally evolved adenine base editors generated through mutation reversion analysis. Nat. Biotechnol. 10.1038/s41587-026-03045-z.

25. Listgarten, J., and Jiang, H. (2026). How artificial intelligence is reengineering protein engineering.

26. Xiao, Y.-L., Wu, Y., and Tang, W. (2024). An adenine base editor variant expands context compatibility. Nature Biotechnology 2024, 1–12. 10.1038/s41587-023-01994-3.

27. Lapinaite, A., Knott, G.J., Palumbo, C.M., Lin-Shiao, E., Richter, M.F., Zhao, K.T., Beal, P.A., Liu, D.R., and Doudna, J.A. (2020). DNA capture by a CRISPR-Cas9– guided adenine base editor. Science (1979). 369, 566–571. 10.1126/science.abb1390.

28. Doman, J.L., Raguram, A., Newby, G.A., and Liu, D.R. (2020). Evaluation and minimization of Cas9-independent off-target DNA editing by cytosine base editors. Nat. Biotechnol. 38, 620–628. 10.1038/s41587-020-0414-6.

29. Grünewald, J., Zhou, R., Garcia, S.P., Iyer, S., Lareau, C.A., Aryee, M.J., and Joung, J.K. (2019). Transcriptome-wide off-target RNA editing induced by CRISPR-guided DNA base editors. Preprint at Nature Publishing Group, http://doi.org/10.1038/s41586-019-1161-z 10.1038/s41586-019-1161-z.

30. Kim, D., Kim, D. eun, Lee, G., Cho, S.I., and Kim, J.S. (2019). Genome-wide target specificity of CRISPR RNA-guided adenine base editors. Nat. Biotechnol. 37, 430–435. 10.1038/s41587-019-0050-1.

31. Tálas, A., Simon, D.A., Kulcsár, P.I., Varga, É., Krausz, S.L., and Welker, E. (2021). BEAR reveals that increased fidelity variants can successfully reduce the mismatch tolerance of adenine but not cytosine base editors. Nat. Commun. 12, 1–14. 10.1038/s41467-021-26461-y.

32. Zhang, Z., Tao, W., Huang, S., Sun, W., Wang, Y., Jiang, W., Huang, X., and Lin, C.P. (2022). Engineering an adenine base editor in human embryonic stem cells with minimal DNA and RNA off-target activities. Mol. Ther. Nucleic Acids 29, 502–510. 10.1016/j.omtn.2022.07.026.

33. Yuan, K., Xi, X., Han, S., Han, J., Zhao, B., Wei, Q., and Zhou, X. (2025). Selict-seq profiles genome-wide off-target effects in adenosine base editing. Nucleic Acids Res. 53. 10.1093/nar/gkaf281.

34. Davis, J.R., Wang, X., Witte, I.P., Huang, T.P., Levy, J.M., Raguram, A., Banskota, S., Seidah, N.G., Musunuru, K., and Liu, D.R. (2022). Efficient in vivo base editing via single adeno-associated viruses with size-optimized genomes encoding compact adenine base editors. Nat. Biomed. Eng. 10.1038/s41551-022-00911-4.

35. Wei, Y., Cao, X., Huang, S., Yue, Y., Luo, M., Liu, Z., Liu, B., Zhang, Q., Wu, Y., Wang, L., et al. (2025). Efficient glycosylase-mediated base editing with minimal off-target effects in mammalian embryos. Genome Biol. 26. 10.1186/s13059-025-03838-6.

36. Cornish-Bowden, A. (1984). Enzyme Specificity: Its Meaning in the General Case. J. Theor. Biol. 108, 451–457.

37. Kim, J., Malashkevich, V., Roday, S., Lisbin, M., Schramm, V.L., and Almo, S.C. (2006). Structural and kinetic characterization of Escherichia coli TadA, the wobble-Specific tRNA deaminase. Biochemistry 45, 6407–6416. 10.1021/bi0522394.

38. Losey, H.C., Ruthenburg, A.J., and Verdine, G.L. (2006). Crystal structure of Staphylococcus aureus tRNA adenosine deaminase TadA in complex with RNA. Nat. Struct. Mol. Biol. 13, 153–159. 10.1038/nsmb1047.

39. Jiang, F., Zhou, K., Ma, L., Gressel, S., and Doudna, J. (2015). A Cas9-guide RNA complex preorganized for target DNA recognition. Science (1979). 348, 1477–1481. 10.1126/science.aab1452.

40. Anders, C., Niewoehner, O., Duerst, A., and Jinek, M. (2014). Structural basis of PAM-dependent target DNA recognition by the Cas9 endonuclease. Nature 513, 569–573. 10.1038/nature13579.

41. Kleinstiver, B.P., Prew, M.S., Tsai, S.Q., Topkar, V. V., Nguyen, N.T., Zheng, Z., Gonzales, A.P.W., Li, Z., Peterson, R.T., Yeh, J.R.J., et al. (2015). Engineered CRISPR-Cas9 nucleases with altered PAM specificities. Nature 523, 481–485. 10.1038/nature14592.

42. Hopf, T.A., Ingraham, J.B., Poelwijk, F.J., Schärfe, C.P.I., Springer, M., Sander, C., and Marks, D.S. (2017). Mutation effects predicted from sequence co-variation. Nat. Biotechnol. 35, 128–135. 10.1038/nbt.3769.

43. Hopf, T.A., Green, A.G., Schubert, B., Mersmann, S., Schärfe, C.P.I., Ingraham, J.B., Toth-Petroczy, A., Brock, K., Riesselman, A.J., Palmedo, P., et al. (2019). The EVcouplings Python framework for coevolutionary sequence analysis. Bioinformatics 35, 1582–1584. 10.1093/bioinformatics/bty862.

44. Eddy, S.R. (2011). Accelerated profile HMM searches. PLoS Comput. Biol. 7. 10.1371/journal.pcbi.1002195.

45. Suzek, B.E., Wang, Y., Huang, H., McGarvey, P.B., and Wu, C.H. (2015). UniRef clusters: A comprehensive and scalable alternative for improving sequence similarity searches. Bioinformatics 31, 926–932. 10.1093/bioinformatics/btu739.

46. Hsu, C., Verkuil, R., Liu, J., Lin, Z., Hie, B., Sercu, T., Lerer, A., and Rives, A. Learning inverse folding from millions of predicted structures.

47. Zhu, D., Brookes, D.H., Busia, A., Carneiro, A., Fannjiang, C., Popova, G., Shin, D., Donohue, K.C., Lin, L.F., Miller, Z.M., et al. (2024). Optimal trade-off control in machine learning-based library design, with application to adeno-associated virus (AAV) for gene therapy.

48. Kunzmann, P., Müller, T.D., Greil, M., Krumbach, J.H., Anter, J.M., Bauer, D., Islam, F., and Hamacher, K. (2023). Biotite: new tools for a versatile Python bioinformatics library. BMC Bioinformatics 24. 10.1186/s12859-023-05345-6.

49. Grathwohl, W., Swersky, K., Hashemi, M., Duvenaud, D., and Maddison, C.J. (2021). Oops I Took A Gradient: Scalable Sampling for Discrete Distributions.

50. Martin, M. Cutadapt removes adapter sequences from high-throughput sequencing reads 10.14806/ej.17.1.200.

51. Pinello, L., Canver, M.C., Hoban, M.D., Orkin, S.H., Kohn, D.B., Bauer, D.E., and Yuan, G.C. (2016). Analyzing CRISPR genome-editing experiments with CRISPResso. Preprint at Nature Publishing Group, http://doi.org/10.1038/nbt.3583 10.1038/nbt.3583.

52. Clement, K., Rees, H., Canver, M.C., Gehrke, J.M., Farouni, R., Hsu, J.Y., Cole, M.A., Liu, D.R., Joung, J.K., Bauer, D.E., et al. (2019). CRISPResso2 provides accurate and reapid genome editing sequence analysis. Preprint at Nature Publishing Group, http://doi.org/10.1038/s41587-019-0046-x 10.1038/s41587-019-0046-x.

53. Picardi, E., and Pesole, G. (2013). REDItools: High-throughput RNA editing detection made easy. Bioinformatics 29, 1813–1814. 10.1093/bioinformatics/btt287.

54. Fonzino, A., Mazzacuva, P.L., Handen, A., Silvestris, D.A., Arnold, A., Pecori, R., Pesole, G., and Picardi, E. (2025). REDInet: a temporal convolutional network-based classifier for A-to-I RNA editing detection harnessing million known events. Brief. Bioinform. 26. 10.1093/bib/bbaf107.

