## Supplementary files for "Activity and specificity trade-offs in adenine base editors"

### Supplementary figures

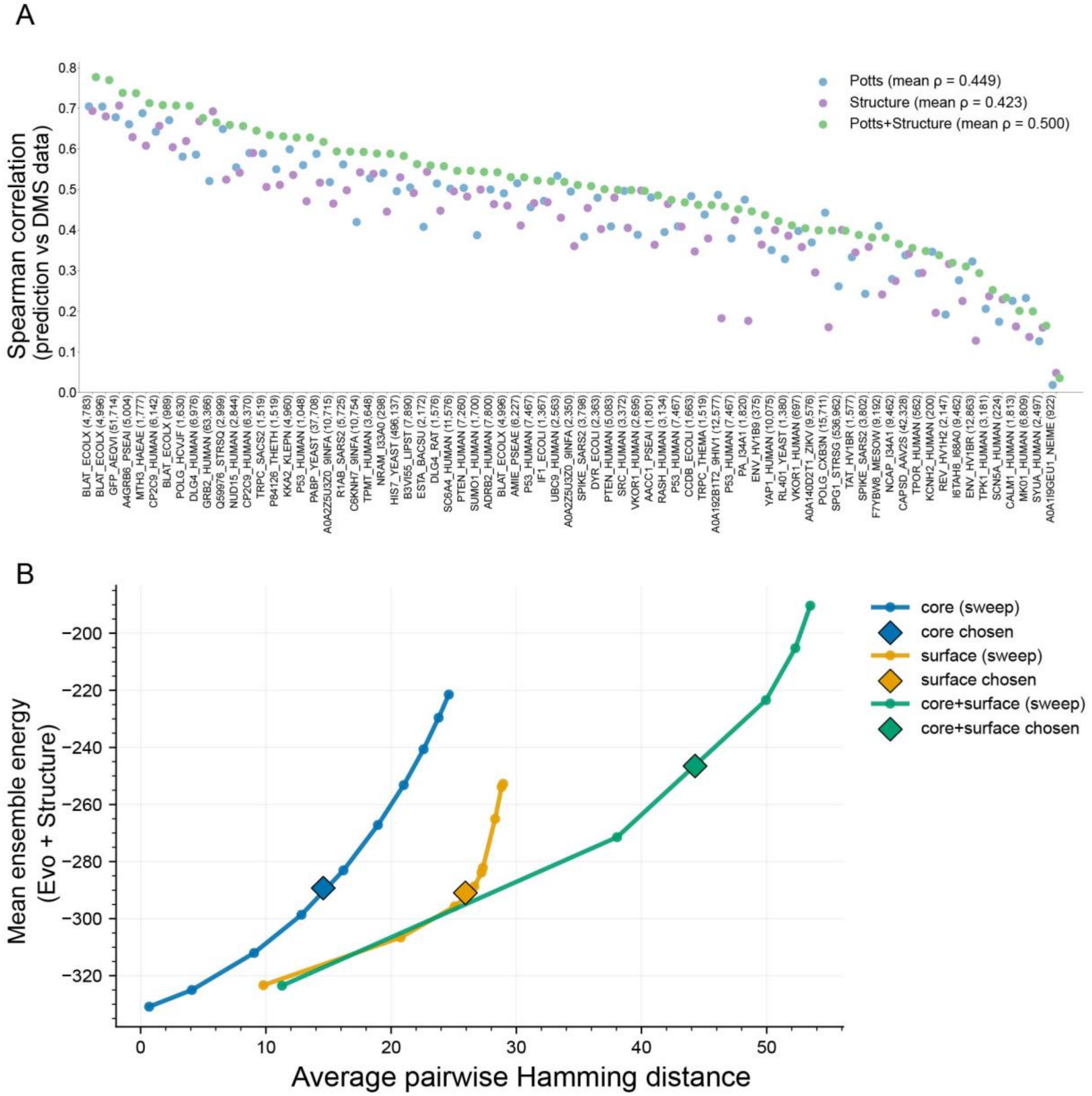

**Figure S1. ML-guided design of ABEs, related to Figure 1.**

A. Ensembling evolutionary and structure models improves zero-shot fitness prediction. Validation of the ensembling strategy used for library design, benchmarked on deep mutational scanning (DMS) datasets from ProteinGym. For each dataset we score every measured variant with two models and compare the predicted scores to the experimental fitness measurements by Spearman correlation (ρ). The evolutionary (Evo) model is trained on a multiple-sequence alignment of homologs and captures sequence constraints and amino-acid covariation; the Structure model is an inverse-folding model (ESM-IF1) distilled into a Potts model, scoring how compatible a sequence is with the protein's structure. The Evo+Structure ensemble is the average of the two per-variant scores. Each point is one dataset (UniProt identifier, with the number of scored variants in parentheses), sorted in descending order of the ensemble correlation (n = 73 datasets). Overall, the ensemble outperforms either model alone (mean ρ = 0.500 vs. 0.449 for Evo and 0.423 for Structure alone), motivating the combined score used to design the ABE library (Figure 1A). B. Trade-off between library diversity and model score in the design procedure. Each ML-guided sublibrary was generated by a simulated-annealing procedure that jointly optimized the ensemble model score and sequence diversity across designs (see Methods). For each of the three sampling strategies, core (blue), surface (orange), and core+surface (green), the diversity weight in the objective was swept to generate a series of candidate libraries. Each circle represents one candidate library, summarized by its average pairwise Hamming distance among designs on the x-axis and its mean ensemble energy on the y-axis. Connected circles of the same color trace the attainable trade-off front for that strategy. Ensemble energy is defined as the negative of the combined Evo + Structure score that guides design, averaged over the designs in each library, so more negative values indicate more favorable designs. Increasing diversity comes at the cost of less favorable scores. Diamonds mark the candidate library chosen for each strategy in the final design, selected to maximize sequence diversity while retaining favorable ensemble scores.

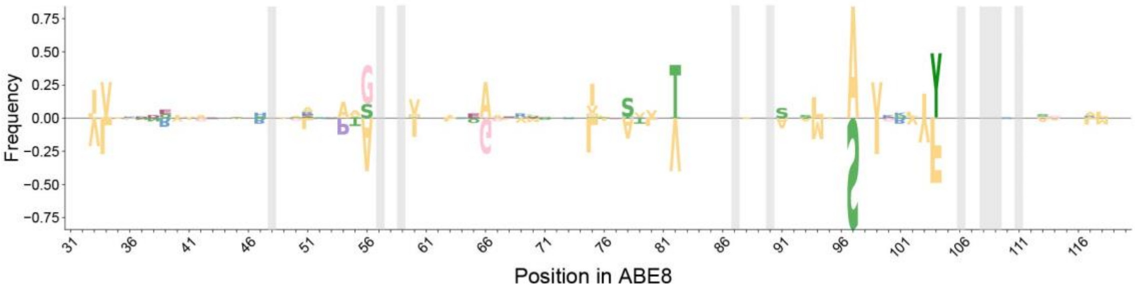

**Figure S2. Logo plot of high activity variants, related to Figure 1.**

Logo plot of the variants with higher than ABE8 activities in the two hard selections (A10 and GA). Fixed residues that are not diversified in the library are indicated with grey rectangles.

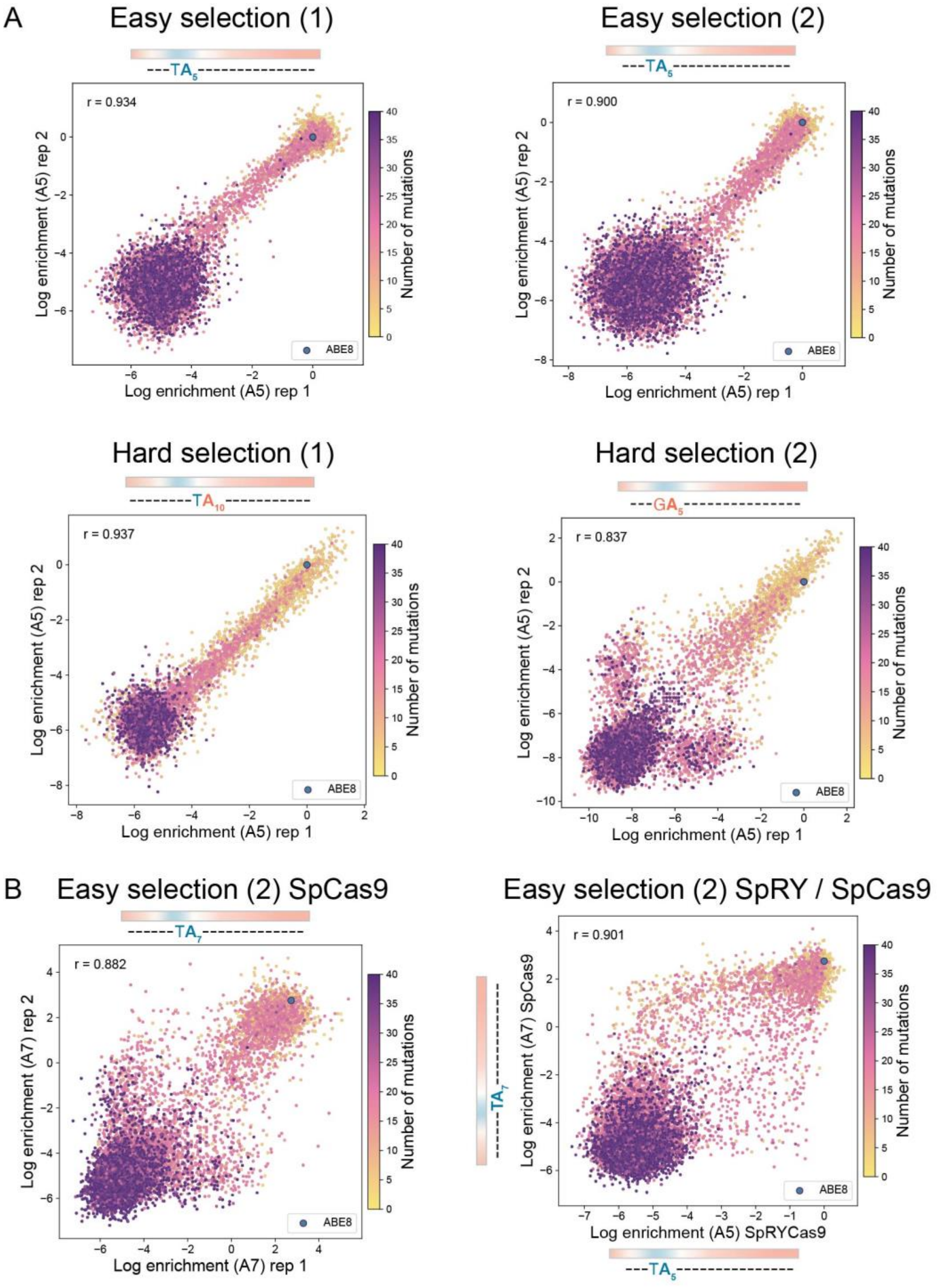

**Figure S3. Correlations between biological replicates of the high-throughput assays and between assays with Sp and SpRY-Cas9-based libraries, related to Figure 1.**

A. Correlations between biological replicates (separate transformations) of the two easy and two hard selection conditions. B. Correlations between replicates of the same library fused to dSpCas9 (left), correlation between the log enrichments of equivalent selections for SpCas9 and SpRY-Cas9 (right). The SpCas9 spacer was adjusted to have a NGG PAM, therefore the target

A was at A7. For the easy SpRY-Cas9 selections the easy selections were at A5 (Table S2). Replicate Pearson correlation is shown on the upper left (r).

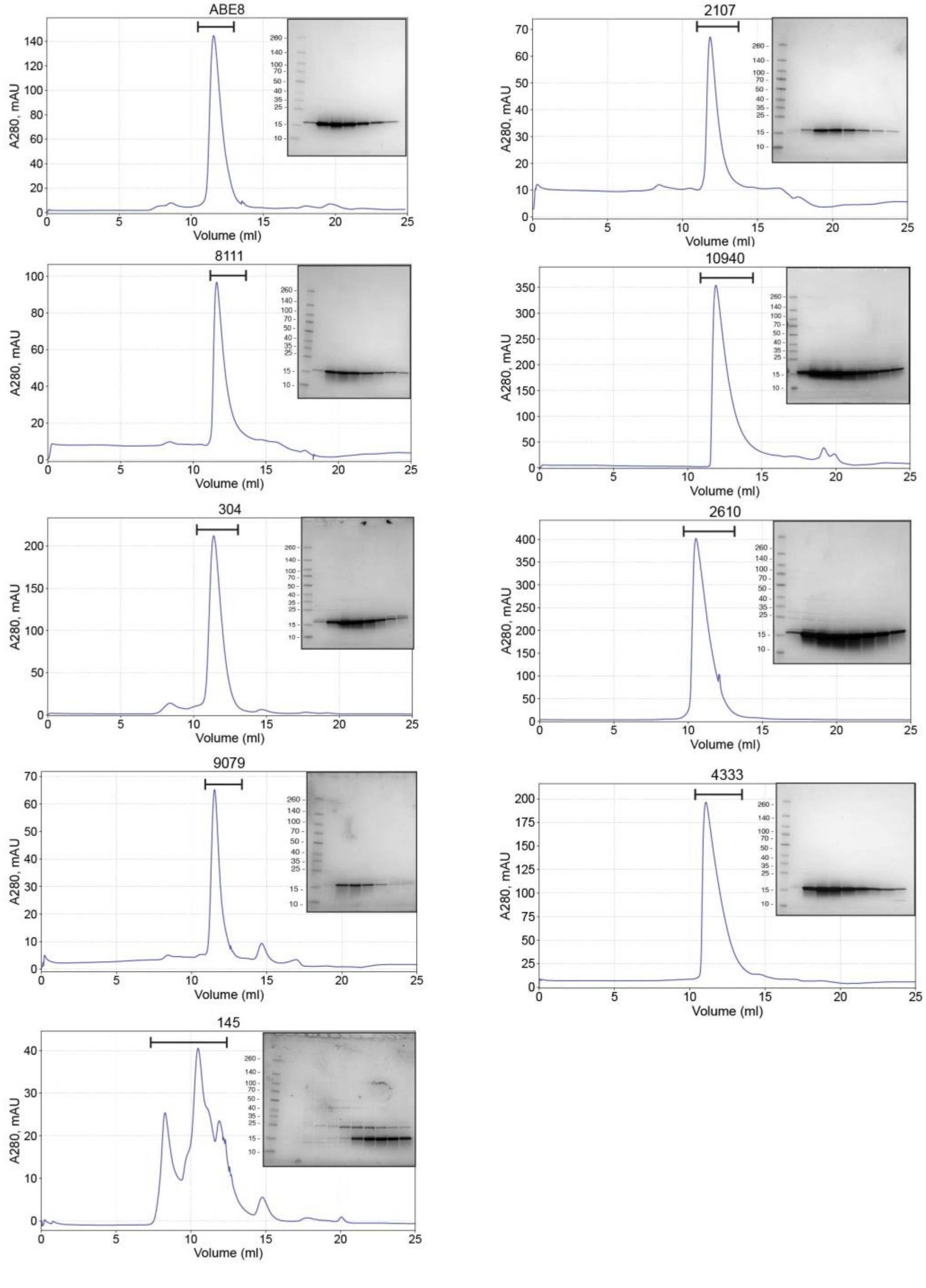

**Figure S4. Purification details of deaminase variants, related to Figure 2.**

Size-exclusion (SEC) profiles of each of the purified variants on a Superdex 75 column. Peak fractions were analyzed by sodium dodecyl sulfate polyacrylamide gel electrophoresis (SDS-PAGE) on a 4-20% gradient gels. The peak concentrated fractions were collected and pooled.

Variant 145 did not have a homogeneous peak and was not pursued. Variant 3925 did not give enough yield after the cation-exchange chromatography and was also not pursued.

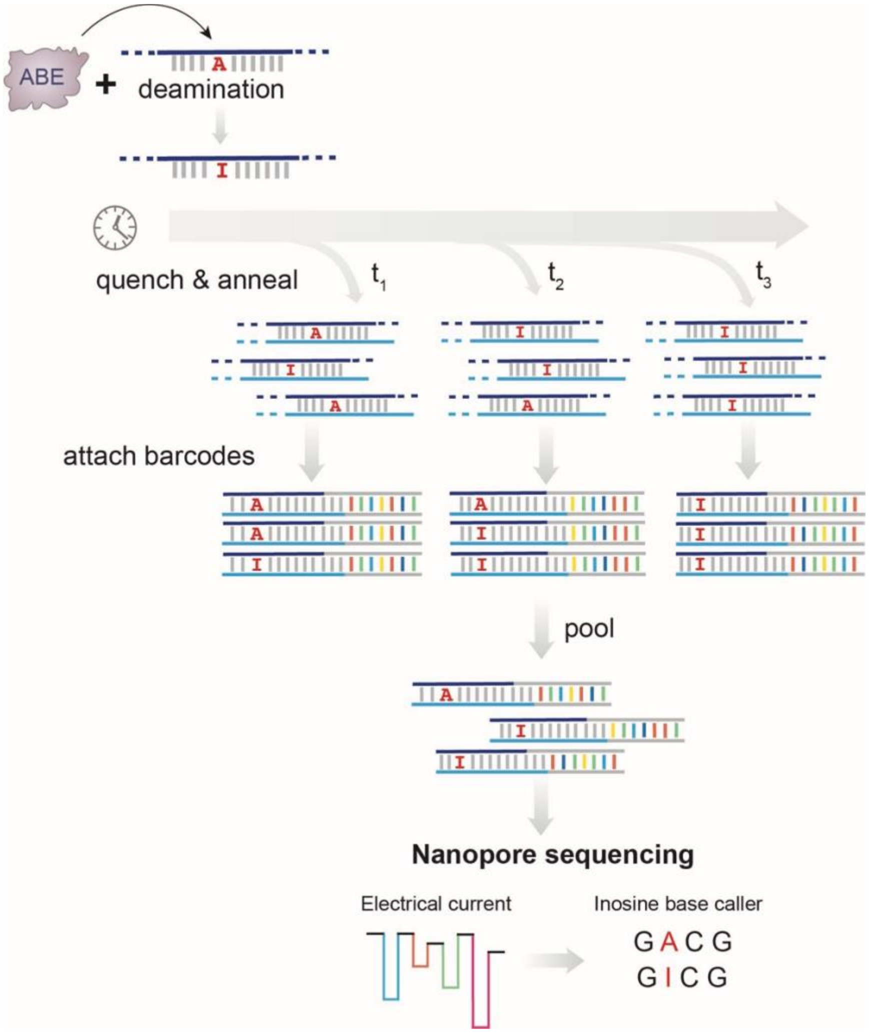

**Figure S5. Schematic of the *in vitro* nanopore-based assay, related to Figure 2.**

Single-stranded DNA oligonucleotide is incubated with the deaminase and at different time points samples are quenched at 95°C. At the same time, a partially complementary strand (with two mismatches against the target A and the preceding nucleotide, see Methods: *In vitro* DNA editing assay with direct inosine detection) are annealed to form a double-stranded (ds) DNA. Each sample is ligated to a unique barcode and samples are pooled and sequenced with nanopore sequencing. The produced inosine can be directly read by the custom inosine base caller model.

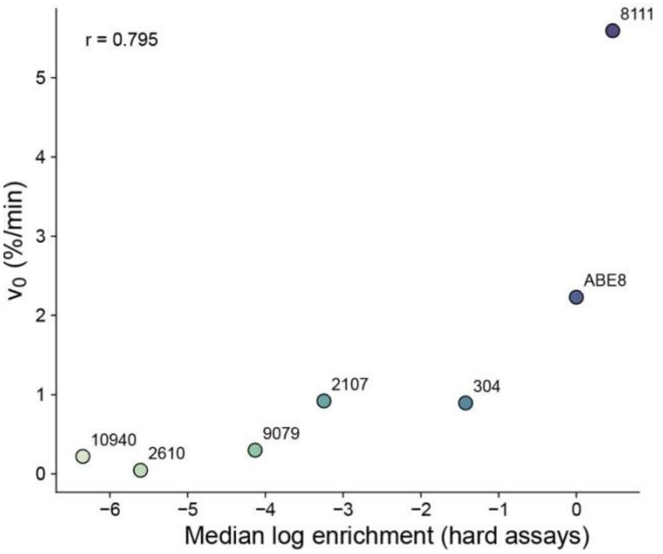

**Figure S6. Correlation between reaction rates and log enrichment measured in the high-throughput assays, related to Figure 2.**

Reaction rates ($V_0$) are calculated as the derived $k_{obs}$ multiplied by the final amplitude reached by each variant (which reflects the active enzyme fraction) in the *in vitro* deamination assay (see Methods: *In vitro* DNA editing assay with direct inosine detection). The rates are plotted against the mean log enrichment from the two hard assays.

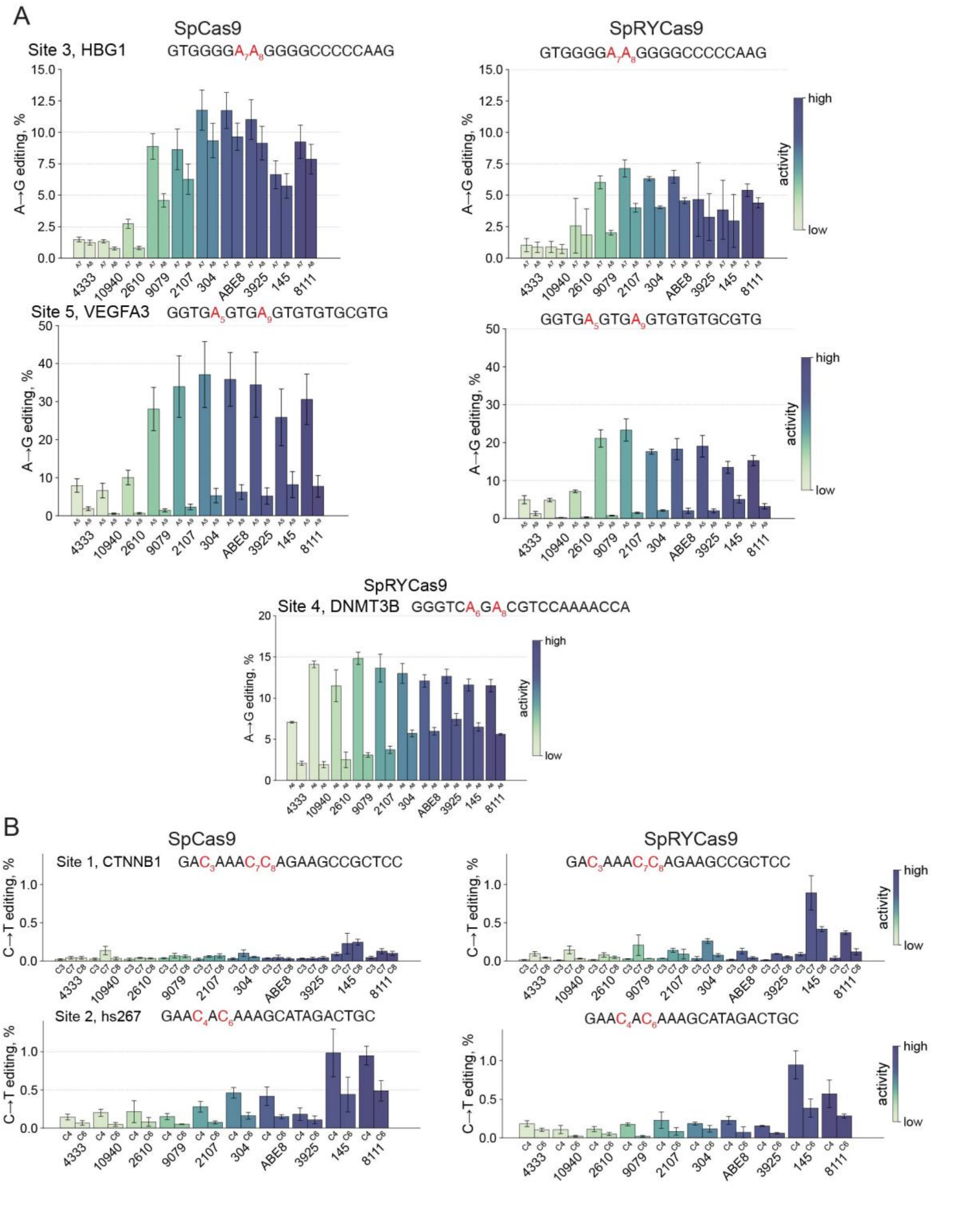

**Figure S7. Editing at endogenous HEK293T sites, related to Figure 2.**

A. Editing at three endogenous HEK293T sites. ABE variants are fused to either nSpCas9 (left panels) or nSpRY-Cas9 (right). Site 4 is only accessible by SpRY-Cas9 due to its PAM. Edited positions shown on the bar graphs are highlighted in red in the sequence. Bars represent mean editing of three biological replicates, each averaged from three technical replicates. Error bars show standard deviation of the biological replicates. B. Cytidine editing within the window of sites 1 and 2 (the A editing is shown on Figure 2C).

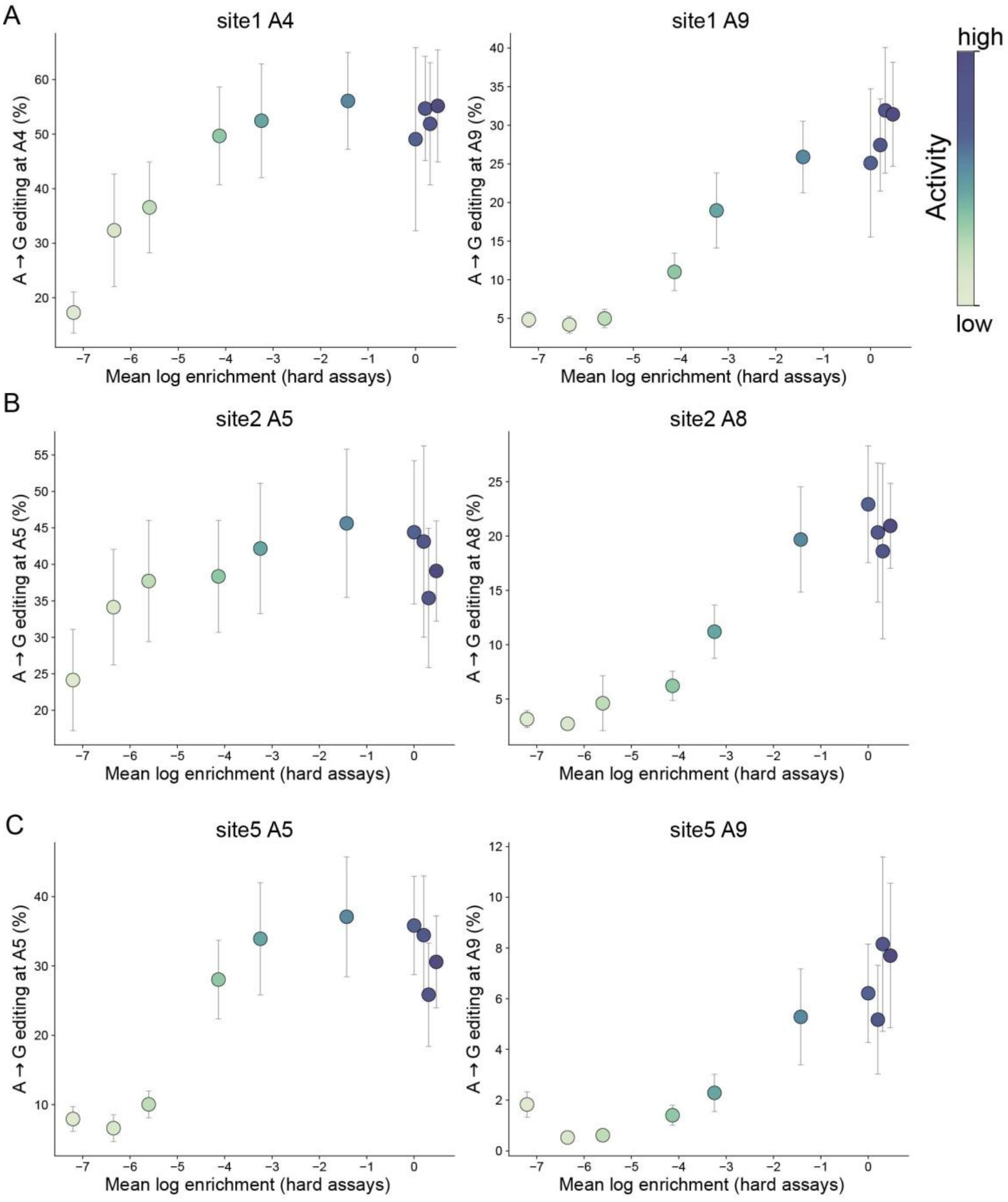

**Figure S8. Editing pattern at three HEK293T sites at middle and the margin of the peak of the editing window, related to Figure 2.**

The editing at the peak and at the margin of the editing window is plotted against the activity (as calculated by the mean log enrichment from the two hard bacterial assays). Variants are also colored according to the same color scheme as on Figure 2A. A. Site 1: position A4 (peak window) and A9 (margin). B. Site 2: position A5 (peak window) and A8 (margin). C. Site 5: position A5 (peak window) and A9 (margin).

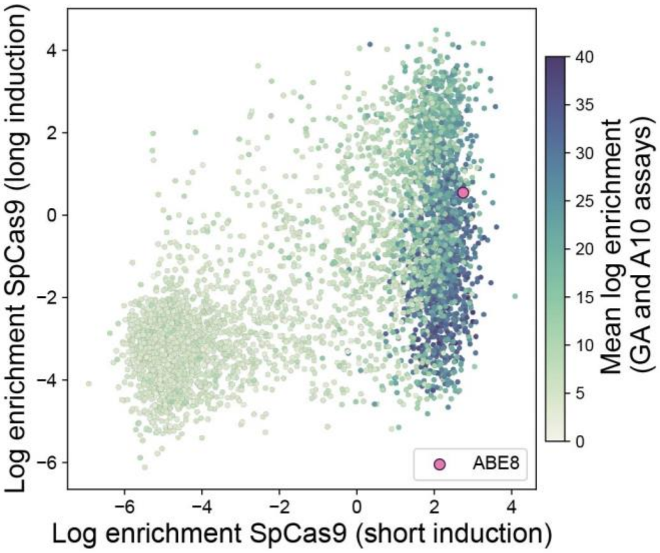

**Figure S9. Toxicity in high-throughput bacterial assay with ABE-SpCas9 library, related to Figure 3.**

The ABE-SpCas9 library was tested in the easy selection (A7) with a short (1h) and long (16h) induction time. Variants are colored by their mean log enrichment for the ABE-SpRY-Cas9 library in the two hard assays (Figure 2A).

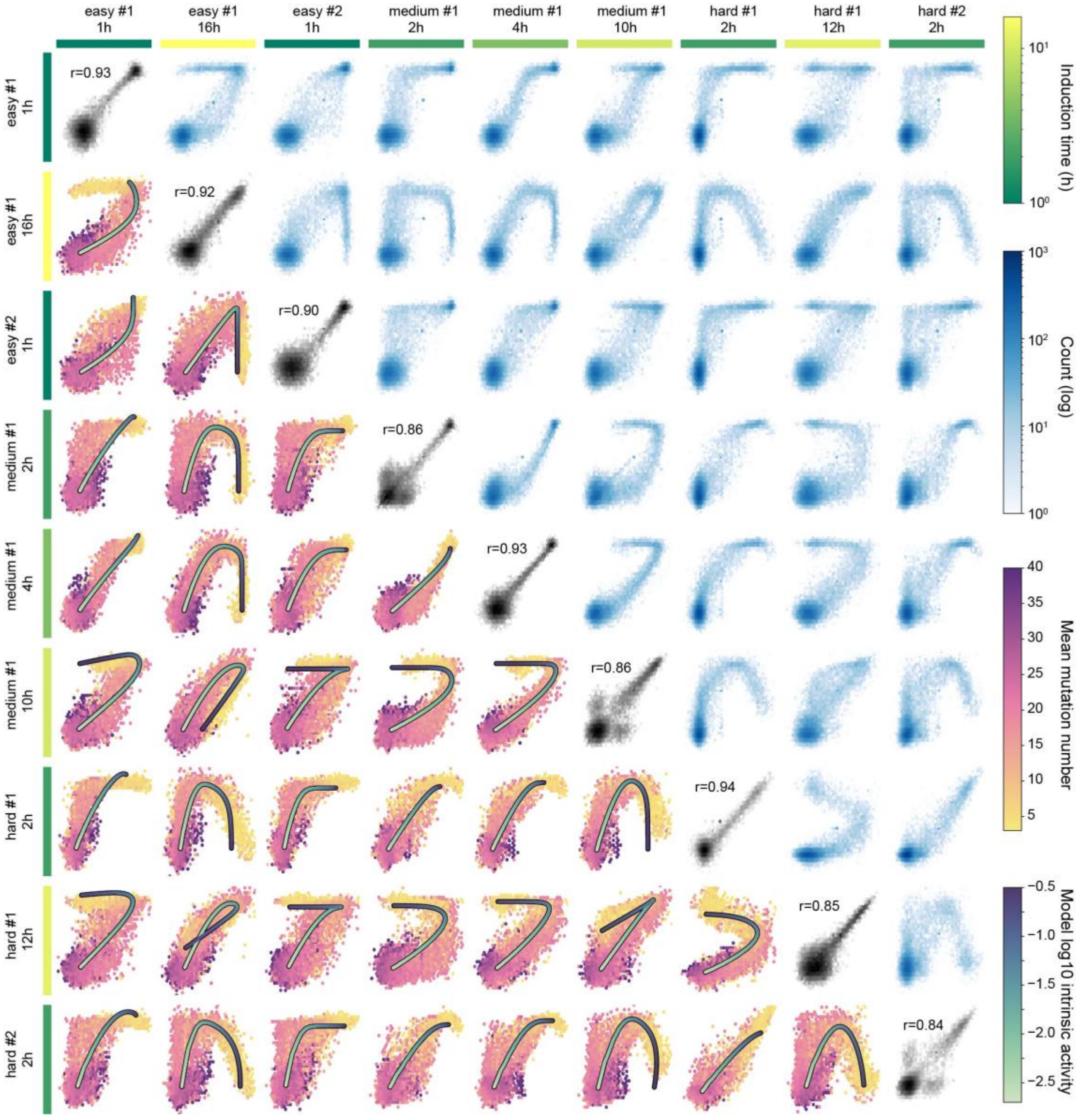

**Figure S10. Toxicity assay, related to Figure 3.**

Arrayed left to right and top to bottom are 9 different assay selection conditions. The labels indicate the particular assay target (e.g. easy #2) and the length of induction time. The induction time is also indicated with the green scale color bar in order to facilitate visual comparison. The upper axis bar color is the induction time of the top assay while the left axis bar color is the left assay induction time.

The plots along the diagonal, i.e. for the same assay with itself, show the variant-by-variant enrichment points for the two replicates. The replicate Pearson correlation is displayed in the upper left of each plot. The upper off-diagonal plots show the pairwise enrichments of the library between different selections. The plots show 2d hexbin histograms in which the log of the counts per bin is according to the blue scale color bar. The key observation is that the density is generally tightly gathered along a one-dimensional manifold in each case. The lower off-diagonal plots are symmetric with the upper off-diagonal (e.g. row 6, column 2 is the same as row 2, column 6). These plots show two types of information. First, the hexbins are colored according to the average number of mutations of the variants populating that bin. Specifically, for hexbins containing 5 or more variants the average number of mutations among those library variants is computed and the bin is colored according to the "mean_n_mutations" color bar (bins with under 5 variants are left white). The second feature of the plots is the parametric curve of the model prediction. Along these curves the intrinsic activity is indicated by the "model log10 intrinsic activity" color bar. These curves were generating by fitting the model parameters in the following way: the off-target activity constant, the cured plasmid survival threshold, and the minimum and maximum intrinsic deaminase activities were all fit single global values. The on-target activity constants were fit to values matching the 4 unique target sequences (i.e. easy #1 & #2, medium #1, and hard #1 & #2). The effective outgrowth time and an enrichment scaling factor were fit to each of the 9 unique assays (i.e. target and induction time combinations). The parametric curves of the simple activity/toxicity model achieve a high degree of qualitative correspondence over the wide range of complex enrichment point manifolds. Furthermore, the average mutation number gradient is clearly apparent along the manifolds in such a way as that lower average mutations localize near the higher activity portions of the parametric curves while high average mutations localize the lower activity positions.

Taken together, we interpret that this collection of data and fits as pointing strongly in favor of a single degree of freedom among the library variants that causes them to cluster along a one-dimensional manifold in a way that is ordered with respect to average mutation number. Moreover, informed by our model, we contend that this single degree of freedom is the intrinsic deaminase activity.

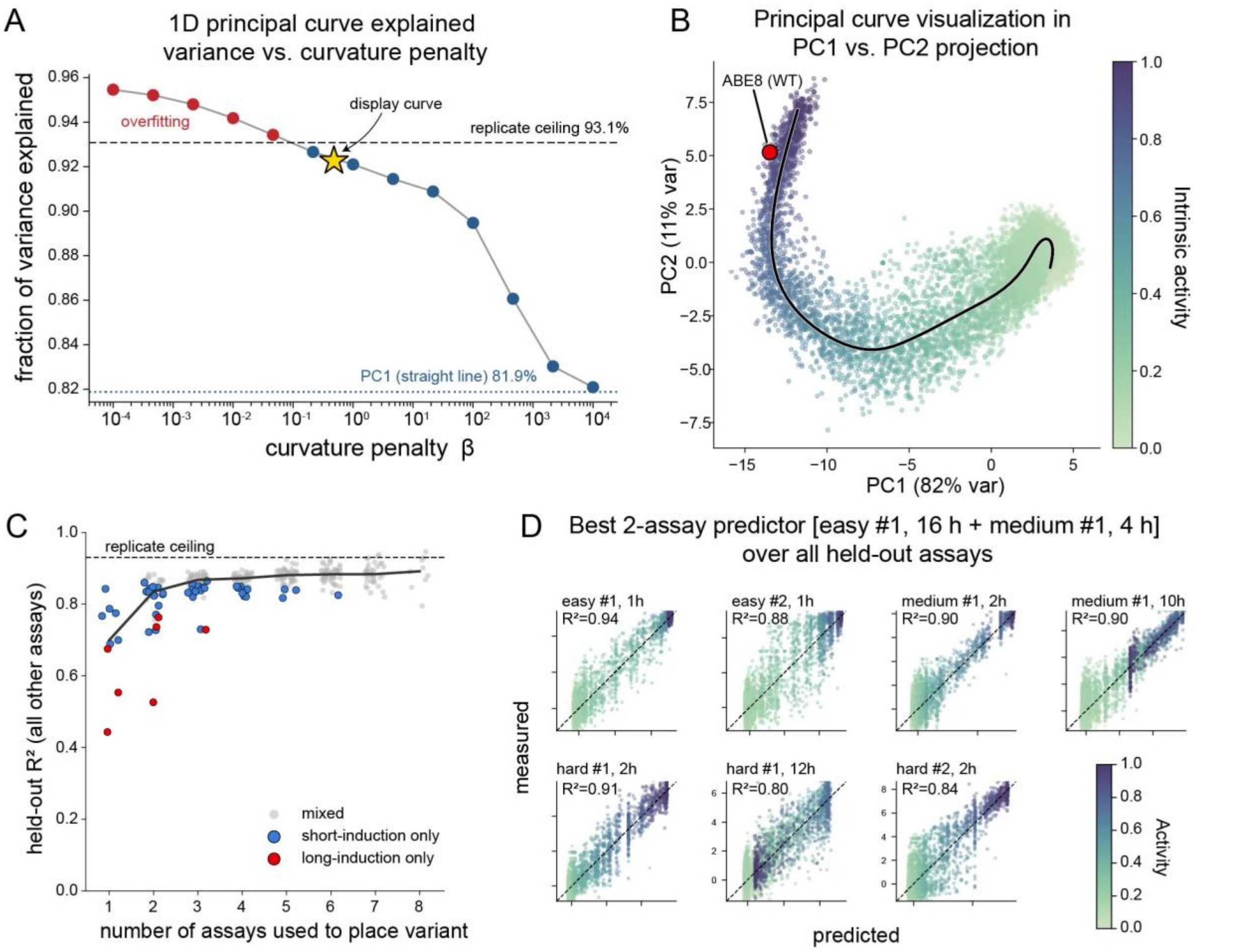

**Figure S11. Model-independent dimensionality analysis, related to Figure 3**

A principal curve was fit to the 9-dimensional library assay data to test the hypothesis that variant behavior is inherently described by a single dimension, independently of the biophysical model. A. Fraction of variance explained (FVE) by the principal curve as a function of the curvature penalty β; larger β forces the curve toward a straight line. Across a broad range of penalties the FVE sits just beneath the replicate-defined reproducibility ceiling — capturing nearly all of the variance that is explainable in principle — and falls toward the linear (PCA) value only at very large β, showing that the one-dimensional description is robust to the choice of penalty. B. The principal curve projected into the plane of the first two linear principal components, to visualize the fit; the β value and FVE are those marked by the star in (A). Points are colored by the fitted activity coordinate, with ABE8 (WT) indicated. C. Held-out validation of the curve's ability to generalize: each point is the $R^2$ for predicting a variant's enrichment in the assays withheld from the fit, as a function of how many assays were revealed to place the variant. The accuracy rapidly plateaus with the number of unmasked assays (within about 3), indicating that a few assays suffice to fix the activity coordinate and that the remaining assays are redundant rather than independent dimensions. Points are colored by subset composition (gray, mixed; blue, short-induction only; red, long-induction only); long-induction-only subsets predict poorly because they sample only the high-activity turnover regime. D. Predicted versus measured enrichment for the best-performing pair of assays (easy #1, 16 h + medium #1, 4 h — one long- and one short-induction condition), used to predict each of the seven remaining masked assays; points colored by activity.

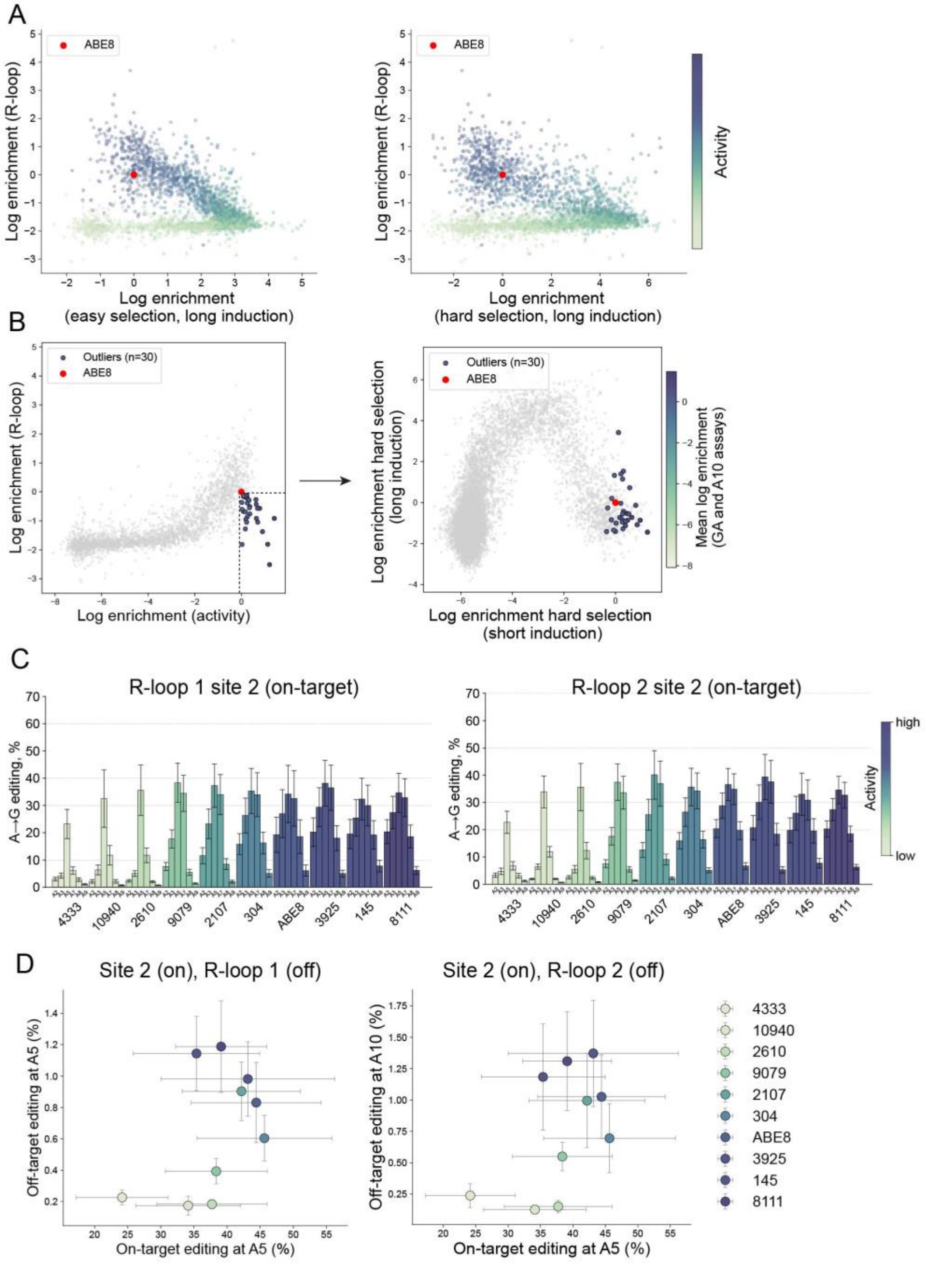

**Figure S12. Bacterial and mammalian orthogonal R-loop assays to measure off-target activity, related to Figure 4.**

A. Log enrichments of the R-loop high-throughput assay and the long induction activity assays, easy (left) and hard (right) (from Figure 3B). B. Variants that have high activity and low R-loop activity (bottom right corner) are highlighted in blue and plotted on the long induction assays (hard target A10), showing that they do have decreased enrichments in the long induction conditions and are therefore not true outliers in all assays. C. Editing at the on-target site for the R-loop off-target assay shown in Figure 4C. The SaCas9 guide targets R-loop site 1 and site 2 shown on Figure 4C and the SpCas9 guide is targeting site 2 (same as site 2 shown on Figure 2C). Bars represent mean editing of three biological replicates, each averaged from three technical replicates. Error bars show standard deviation of the biological replicates. D. Off-target R-loop editing (Figure 4C) is plotted against the on-target editing (C) in HEK293T cells, showing the saturation of the on-target editing, while the off-target editing keeps increasing with the deaminase activity.

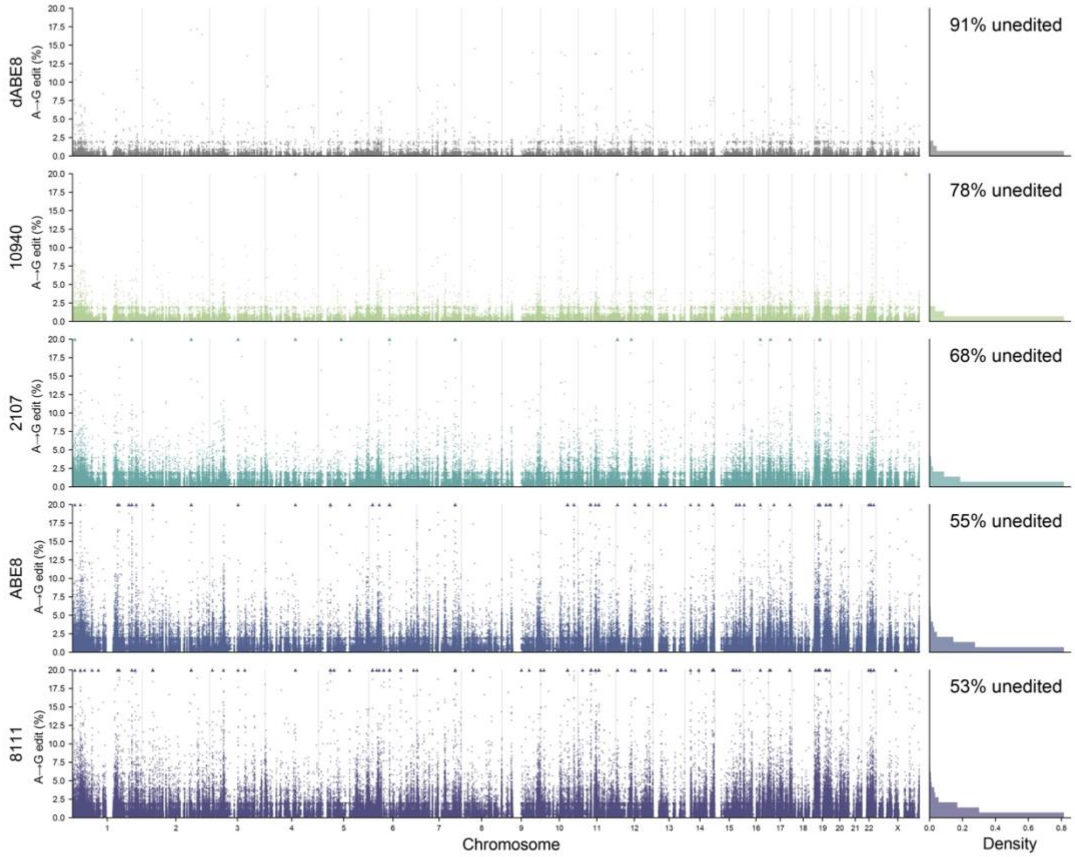

**Figure S13. Transcriptome-wide RNA editing by genomic positions in HEK293T cells measured by RNAseq, related to Figure 4.**

Genome coordinate plot of the A to G editing efficiency at all A's occurring with a read depth of at least 50 for each of the tested variants. Positions with 20% or greater are displayed as triangles and clipped to 20%. Along the right margin are normalized histograms of the editing percentages. Sites with no editing are not included in the histogram but are indicated as the annotated % unedited. Sites blacklisted as putative SNPs are not included in the analysis.

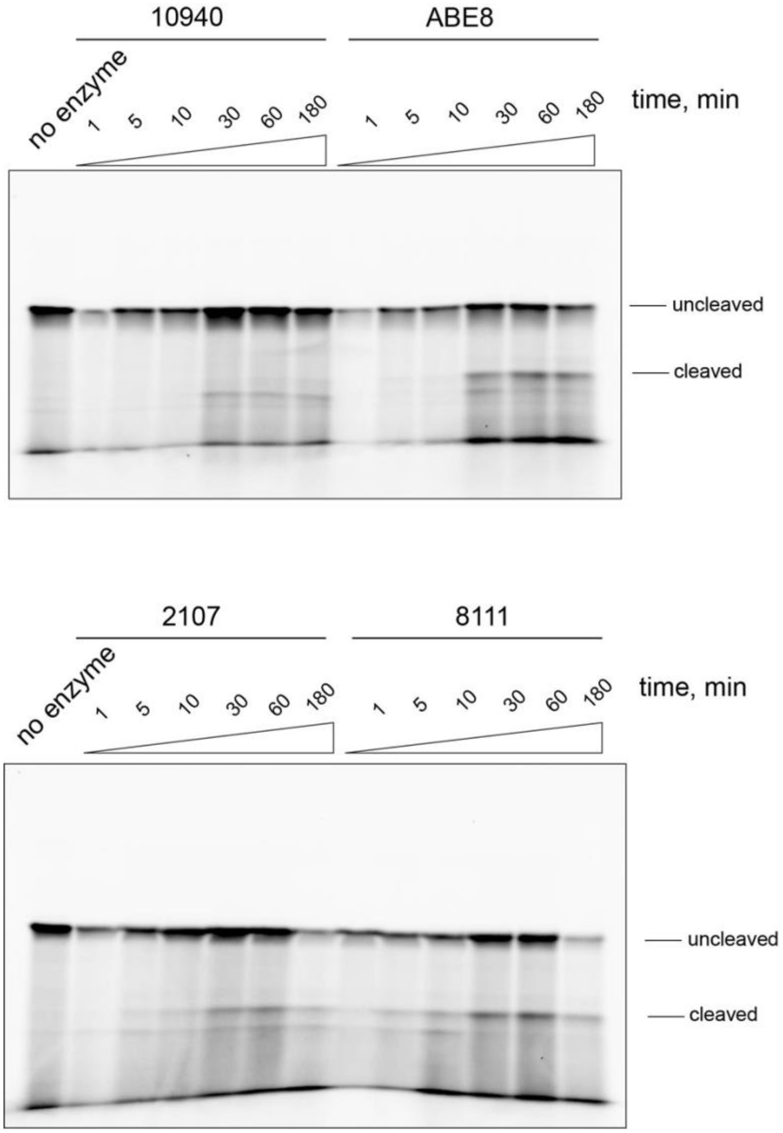

**Figure S14. Representative gels of in vitro RNA editing reactions, related to Figure 4.**

RNA editing was measured by EndoV-coupled reactions, where the RNA containing inosine is cleaved specifically by EndoV endonuclease leading to a cleavage product that can be observed on denaturing polyacrylamide gel (see Methods: *In vitro* RNA editing assay). One typical gel is shown per variant, three separate reactions were used for quantification of the bands to produce the graph on Figure 4E.

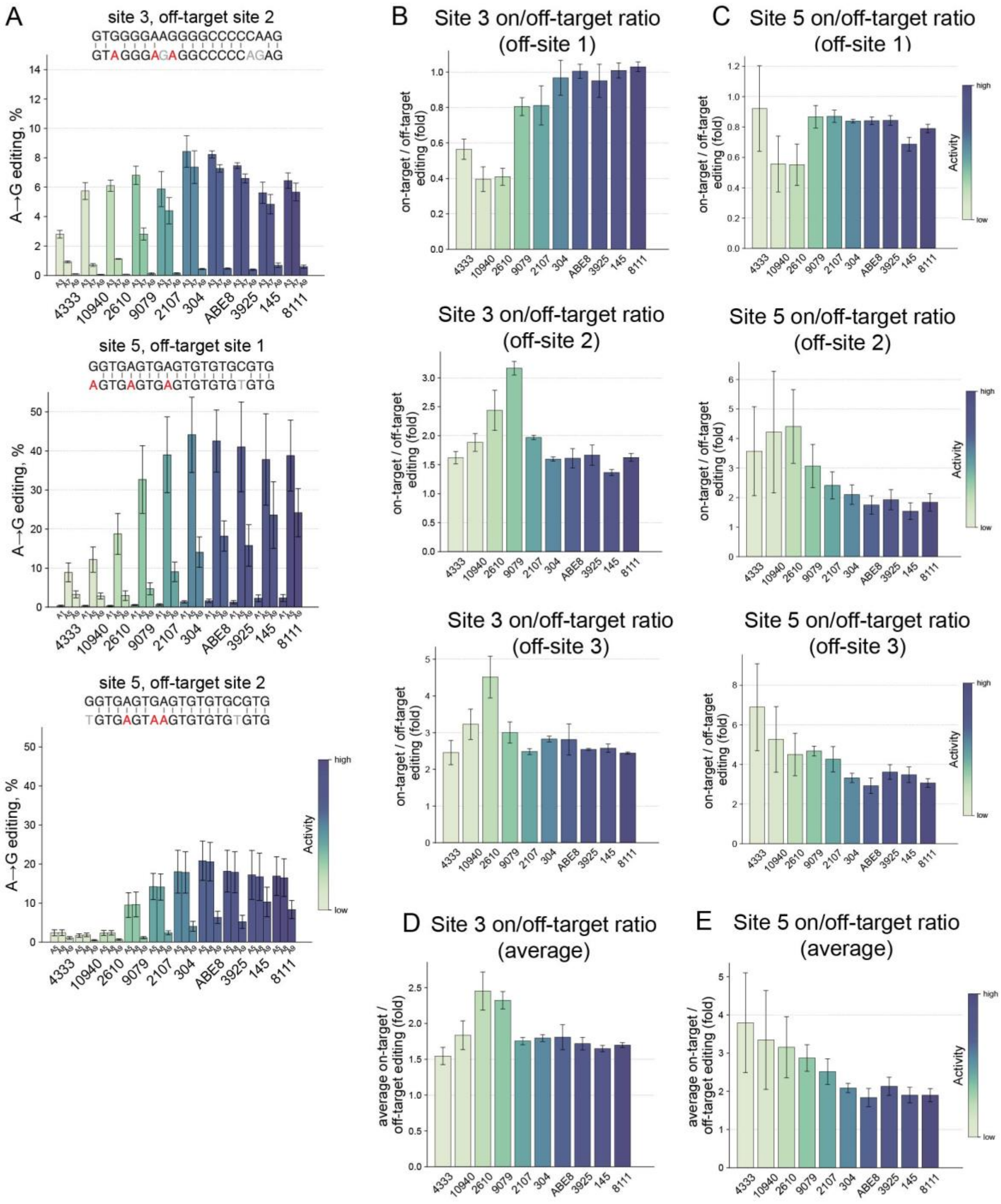

**Figure S15. Cas9-dependent off-targets, related to Figure 5.**

A. Bystander editing at known Cas9-dependent off-target sites. Editing at all positions for the targets shown on Figure 5A. Site 3 off-target site 2 is the only one that contains more than one A. Bars indicate mean editing from three biological replicates, each averaged from three technical replicates, error bars represent standard deviations. Off-target sites 1 and 2 for site 5 have more than one A. B, C. Ratio of on-target to off-target editing for each individual off-target site of site 3 (B), site 5 (C). D, E. Averaged on-target to off-target ratio for all measured off-target sites of site 3 (D) and site 5 (E).

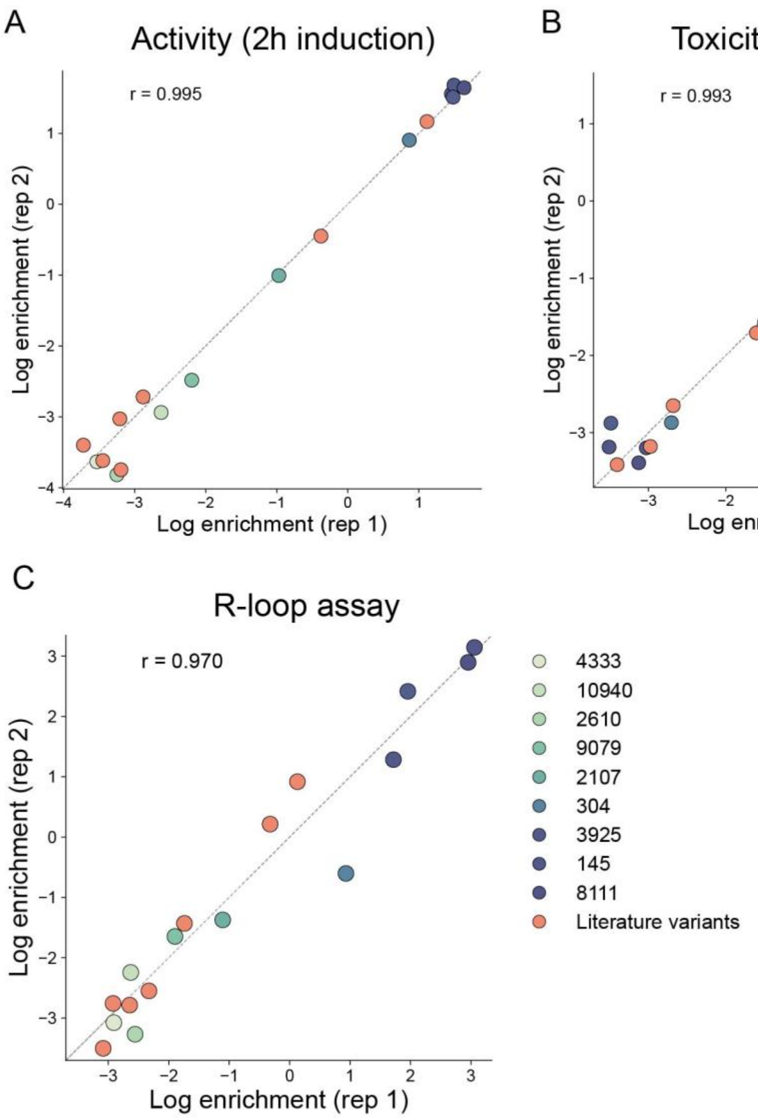

**Figure S16. Replicate correlations of bacterial assays of the subset of library variants and literature variants, related to Figure 6.**

Correlations between biological replicates of the assays testing literature variants together with the subset of library variants with known activities. Literature variants are colored in orange and the library variants are colored according to their activity as show in Figure 2A. A. Log enrichment correlation of replicates in hard selection #1 (A10), short induction (2h) B. Log enrichment correlations of hard selection #1 (A10), long induction (16h). C. Log enrichment correlations of orthogonal R-loop assay for off-target detection. Pearson correlations between the replicates are shown on each graph.

### Supplementary tables

#### Table S1. Fixed residues in the library

Residues that were kept constant because they were either part of the active site or have been shown to be important for the ABE7 or ABE8 evolution. E.g., S48A mutation boosted the activity of ABE6 to ABE7; V106 and N108 were the first residues that enabled DNA editing in the evolution of TadA [1]. ABE8 evolution: R111 reversion back to T leads to significant reduction in activity [2].

| Fixed residues | Motivation |
|---|---|
| A48 | ABE7.10 |
| H57 | Active site |
| E59 | Active site |
| C87 | Active site |
| C90 | Active site |
| V106 | ABE7.10 |
| N108 | ABE7.10 |
| S109 | ABE8 |
| R111 | ABE8 |

**Table S2. Spacers and guides for the bacterial high-throughput selections**

**On-target selections**

| Selection | Spacer 5'-3' | Guide |
|---|---|---|
| Easy selection #1 | AACTAATCTTCAGCATCTTT | AACTAATCTTCAGCATCTTT |
| Easy selection #2 | TCTTACCCGGCGTCAATACG | TCTTACCCGGCGTCAATACG |
| Easy selection #2 (Sp) | GCTCTTACCCGGCGTCAATA | GCTCTTACCCGGCGTCAATA |
| Hard selection #1 | GTTGCTCTTACCCGGCGTCA | GTTGCTCTTACCCGGCGTCA |
| Hard selection #2 | AGTGAGGTACATCGAACTGG | AGTGAGGTACATCGAACTGG |

**Off-target selections**

| | | |
|---|---|---|
| R-loop | TTGCTCTTACCCGGCGTCAATA | TTGCTCTTACCCGGCGTCAATA |
| Mismatched selection (3-5) | AACTAATCTTCAGCATCTTT | AAGCGATCTTCAGCATCTTT |
| Mismatched selection (6-8) | AACTAATCTTCAGCATCTTT | AACTACGATTCAGCATCTTT |
| Mismatched selection (9-11) | AACTAATCTTCAGCATCTTT | AACTAATCGCAAGCATCTTT |

**Table S3. Incubation times for bacterial selections**

**On-target selections**

| Selection | Short induction | Long induction |
|---|---|---|
| Easy selection #1 | 1h | 12h |
| Easy selection #2 | 1h | - |
| Easy selection #2 (Sp) | 1h | 16h |
| Hard selection #1 | 2h | 12h |
| Hard selection #2 | 2h | - |

**Off-target selections**

| | |
|---|---|
| R-loop | 8h |
| Mismatched selection (3-5) | 1h |
| Mismatched selection (6-8) | 1h |
| Mismatched selection (9-11) | 1h |

### Supplementary sequences

#### Reporters

##### Easy selection #1 (A5)

ATGAGTATTCAACATTTCCGTGTCGCCCTTATTCCCTTTTTTGCGGCATTTTGCCTTCCTGTT
TTTGCTCACCCAGAAACGCTGGTGAAAGTA**AAAGATGCTGAAGATTAGTT**GGGTGCACGA
GTGGGTTACATCGAACTGGATCTCAACAGCGGTAAGATCCTTGAGAGTTTTCGCCCCGAAG
AACGTTTTCCAATGATGAGCACTTTTAAAGTTCTGCTATGTGGCGCGGTATTATCCCGTATTG
ACGCCGGGCAAGAGCAACTCGGTCGCCGCATACACTATTCTCAGAATGACTTGGTTGAGTA
CTCACCAGTCACAGAAAAGCATCTTACGGATGGCATGACAGTAAGAGAATTATGCAGTGCTG
CCATAACCATGAGTGATAACACTGCGGCCAACTTACTTCTGACAACGATCGGAGGACCGAA
GGAGCTAACCGCTTTTTTGCACAACATGGGGGATCATGTAACTCGCCTTGATCGTTGGGAA
CCGGAGCTGAATGAAGCCATACCAAACGACGAGCGTGACACCACGATGCCTGTAGCAATG
GCAACAACGTTGCGCAAACTATTAACTGGCGAACTACTTACTCTAGCTTCCCGGCAACAATT
AATAGACTGGATGGAGGCGGATAAAGTTGCAGGACCACTTCTGCGCTCGGCCCTTCCGGC
TGGCTGGTTTATTGCTGATAAATCTGGAGCCGGTGAGCGTGGGTCTCGCGGTATCATTGCA
GCACTGGGGCCAGATGGTAAGCCCTCCCGTATCGTAGTTATCTACACGACGGGGAGTCAGG
CAACTATGGATGAACGAAATAGACAGATCGCTGAGATAGGTGCCTCACTGATTAAGCATTGG
TAA

##### Easy selection #2 (A5)

ATGAGTATTCAACATTTCCGTGTCGCCCTTATTCCCTTTTTTGCGGCATTTTGCCTTCCTGTT
TTTGCTCACCCAGAAACGCTGGTGAAAGTAAAAGATGCTGAAGATCAGTTGGGTGCACGAG
TGGGTTACATCGAACTGGATCTCAACAGCGGTAAGATCCTTGAGAGTTTTCGCCCCGAAGA
ACGTTTTCCAATGATGAGCACTTTTAAAGTTCTGCTATGTGGCGCGGTATTATCC**CGTATTGA
CGCCGGGTAAGA**GCAACTCGGTCGCCGCATACACTATTCTCAGAATGACTTGGTTGAGTAC
TCACCAGTCACAGAAAAGCATCTTACGGATGGCATGACAGTAAGAGAATTATGCAGTGCTGC
CATAACCATGAGTGATAACACTGCGGCCAACTTACTTCTGACAACGATCGGAGGACCGAAG
GAGCTAACCGCTTTTTTGCACAACATGGGGGATCATGTAACTCGCCTTGATCGTTGGGAAC
CGGAGCTGAATGAAGCCATACCAAACGACGAGCGTGACACCACGATGCCTGTAGCAATGG
CAACAACGTTGCGCAAACTATTAACTGGCGAACTACTTACTCTAGCTTCCCGGCAACAATTA
ATAGACTGGATGGAGGCGGATAAAGTTGCAGGACCACTTCTGCGCTCGGCCCTTCCGGCT
GGCTGGTTTATTGCTGATAAATCTGGAGCCGGTGAGCGTGGGTCTCGCGGTATCATTGCAG
CACTGGGGCCAGATGGTAAGCCCTCCCGTATCGTAGTTATCTACACGACGGGGAGTCAGGC
AACTATGGATGAACGAAATAGACAGATCGCTGAGATAGGTGCCTCACTGATTAAGCATTGGT
AA

##### Easy selection #2 (Sp) (A7)

ATGAGTATTCAACATTTCCGTGTCGCCCTTATTCCCTTTTTTGCGGCATTTTGCCTTCCTGTT
TTTGCTCACCCAGAAACGCTGGTGAAAGTAAAAGATGCTGAAGATCAGTTGGGTGCACGAG
TGGGTTACATCGAACTGGATCTCAACAGCGGTAAGATCCTTGAGAGTTTTCGCCCCGAAGA
ACGTTTTCCAATGATGAGCACTTTTAAAGTTCTGCTATGTGGCGCGGTATTATCCCG**TATTGA
CGCCGGGTAAGAGC**AACTCGGTCGCCGCATACACTATTCTCAGAATGACTTGGTTGAGTAC
TCACCAGTCACAGAAAAGCATCTTACGGATGGCATGACAGTAAGAGAATTATGCAGTGCTGC
CATAACCATGAGTGATAACACTGCGGCCAACTTACTTCTGACAACGATCGGAGGACCGAAG
GAGCTAACCGCTTTTTTGCACAACATGGGGGATCATGTAACTCGCCTTGATCGTTGGGAAC
CGGAGCTGAATGAAGCCATACCAAACGACGAGCGTGACACCACGATGCCTGTAGCAATGG

CAACAACGTTGCGCAAACTATTAACTGGCGAACTACTTACTCTAGCTTCCCGGCAACAATTA
ATAGACTGGATGGAGGCGGATAAAGTTGCAGGACCACTTCTGCGCTCGGCCCTTCCGGCT
GGCTGGTTTATTGCTGATAAATCTGGAGCCGGTGAGCGTGGGTCTCGCGGTATCATTGCAG
CACTGGGGCCAGATGGTAAGCCCTCCCGTATCGTAGTTATCTACACGACGGGGAGTCAGGC
AACTATGGATGAACGAAATAGACAGATCGCTGAGATAGGTGCCTCACTGATTAAGCATTGGT
AA

**Hard selection #1 (A10)**

ATGAGTATTCAACATTTCCGTGTCGCCCTTATTCCCTTTTTTGCGGCATTTTGCCTTCCTGTT
TTTGCTCACCCAGAAACGCTGGTGAAAGTAAAAGATGCTGAAGATCAGTTGGGTGCACGAG
TGGGTTACATCGAACTGGATCTCAACAGCGGTAAGATCCTTGAGAGTTTTCGCCCCGAAGA
ACGTTTTCCAATGATGAGCACTTTTAAAGTTCTGCTATGTGGCGCGGTATTATCCCGTAT**TGA**
**CGCCGGGTAAGAGCAAC**TCGGTCGCCGCATACACTATTCTCAGAATGACTTGGTTGAGTAC
TCACCAGTCACAGAAAAGCATCTTACGGATGGCATGACAGTAAGAGAATTATGCAGTGCTGC
CATAACCATGAGTGATAACACTGCGGCCAACTTACTTCTGACAACGATCGGAGGACCGAAG
GAGCTAACCGCTTTTTTGCACAACATGGGGGATCATGTAACTCGCCTTGATCGTTGGGAAC
CGGAGCTGAATGAAGCCATACCAAACGACGAGCGTGACACCACGATGCCTGTAGCAATGG
CAACAACGTTGCGCAAACTATTAACTGGCGAACTACTTACTCTAGCTTCCCGGCAACAATTA
ATAGACTGGATGGAGGCGGATAAAGTTGCAGGACCACTTCTGCGCTCGGCCCTTCCGGCT
GGCTGGTTTATTGCTGATAAATCTGGAGCCGGTGAGCGTGGGTCTCGCGGTATCATTGCAG
CACTGGGGCCAGATGGTAAGCCCTCCCGTATCGTAGTTATCTACACGACGGGGAGTCAGGC
AACTATGGATGAACGAAATAGACAGATCGCTGAGATAGGTGCCTCACTGATTAAGCATTGGT
AA

**Hard selection #2 (GA)**

ATGAGTATTCAACATTTCCGTGTCGCCCTTATTCCCTTTTTTGCGGCATTTTGCCTTCCTGTT
TTTGCTCACCCAGAAACGCTGGTGAAAGTAAAAGATGCTGAAGATCAGTTGGGTGCACG**AG**
**TGAGGTACATCGAACTGG**ATCTCAACAGCGGTAAGATCCTTGAGAGTTTTCGCCCCGAAGA
ACGTTTTCCAATGATGAGCACTTTTAAAGTTCTGCTATGTGGCGCGGTATTATCCCGTATTGA
CGCCGGGCAAGAGCAACTCGGTCGCCGCATACACTATTCTCAGAATGACTTGGTTGAGTAC
TCACCAGTCACAGAAAAGCATCTTACGGATGGCATGACAGTAAGAGAATTATGCAGTGCTGC
CATAACCATGAGTGATAACACTGCGGCCAACTTACTTCTGACAACGATCGGAGGACCGAAG
GAGCTAACCGCTTTTTTGCACAACATGGGGGATCATGTAACTCGCCTTGATCGTTGGGAAC
CGGAGCTGAATGAAGCCATACCAAACGACGAGCGTGACACCACGATGCCTGTAGCAATGG
CAACAACGTTGCGCAAACTATTAACTGGCGAACTACTTACTCTAGCTTCCCGGCAACAATTA
ATAGACTGGATGGAGGCGGATAAAGTTGCAGGACCACTTCTGCGCTCGGCCCTTCCGGCT
GGCTGGTTTATTGCTGATAAATCTGGAGCCGGTGAGCGTGGGTCTCGCGGTATCATTGCAG
CACTGGGGCCAGATGGTAAGCCCTCCCGTATCGTAGTTATCTACACGACGGGGAGTCAGGC
AACTATGGATGAACGAAATAGACAGATCGCTGAGATAGGTGCCTCACTGATTAAGCATTGGT
AA

**R-loop selection**

ATGAGTATTCAACATTTCCGTGTCGCCCTTATTCCCTTTTTTGCGGCATTTTGCCTTCCTGTT
TTTGCTCACCCAGAAACGCTGGTGAAAGTAAAAGATGCTGAAGATCAGTTGGGTGCACGAG
TGGGTTACATCGAACTGGATCTCAACAGCGGTAAGATCCTTGAGAGTTTTCGCCCCGAAGA
ACGTTTTCCAATGATGAGCACTTTTAAAGTTCTGCTATGTGGCGCGGTATTATCCCGT**ATTGA**
**CGCCGGGTAAGAGC**AACTCGGTCGCCGCATACACTATTCTCAGAATGACTTGGTTGAGTAC
TCACCAGTCACAGAAAAGCATCTTACGGATGGCATGACAGTAAGAGAATTATGCAGTGCTGC

CATAACCATGAGTGATAACACTGCGGCCAACTTACTTCTGACAACGATCGGAGGACCGAAG
GAGCTAACCGCTTTTTTGCACAACATGGGGGATCATGTAACTCGCCTTGATCGTTGGGAAC
CGGAGCTGAATGAAGCCATACCAAACGACGAGCGTGACACCACGATGCCTGTAGCAATGG
CAACAACGTTGCGCAAACTATTAACTGGCGAACTACTTACTCTAGCTTCCCGGCAACAATTA
ATAGACTGGATGGAGGCGGATAAAGTTGCAGGACCACTTCTGCGCTCGGCCCTTCCGGCT
GGCTGGTTTATTGCTGATAAATCTGGAGCCGGTGAGCGTGGGTCTCGCGGTATCATTGCAG
CACTGGGGCCAGATGGTAAGCCCTCCCGTATCGTAGTTATCTACACGACGGGGAGTCAGGC
AACTATGGATGAACGAAATAGACAGATCGCTGAGATAGGTGCCTCACTGATTAAGCATTGGT
AA

### *In vitro* DNA substrates

ssDNA (60-mer), deamination substrate

5'-
TCTCGCCGCTGTCTGGCGCCGCTCGTCCT**A**CGTGTCTCGTCGGGCGTCGTCTGTGCGCC
G-3'

Complementary DNA (56-mer)

5'-GCACAGACGACGCCCGACGAGACACGAGGGACGAGCGGCGCCAGACAGCGGCGAGA-
3'

Barcodes:

| | |
|---|---|
| barcode1 | CCAGGATCCTAGGCTGCTGCCACACCTCTTCGATAGCAGTTTGAGAAGCACACGGT |
| barcode2 | CCAGGATCCTAGGCTGCTGCTGTGACTCAGCGTAGATTAGTTGAGAAGCACACGGT |
| barcode3 | CCAGGATCCTAGGCTGCTGCCGGCGCCAGGTAGCGTAATCTTGAGAAGCACACGGT |
| barcode4 | CCAGGATCCTAGGCTGCTGCGGCTCTAGCATACTTAGAGCTTGAGAAGCACACGGT |
| barcode5 | CCAGGATCCTAGGCTGCTGCAGAGTCTTATCCTAACTAACTTGAGAAGCACACGGT |
| barcode6 | CCAGGATCCTAGGCTGCTGCTACCGACCCAGACAAGGGCCTTGAGAAGCACACGGT |
| barcode7 | CCAGGATCCTAGGCTGCTGCGGTGTTAGGGCCAGGTTAAGTTGAGAAGCACACGGT |
| barcode8 | CCAGGATCCTAGGCTGCTGCTACTTCCTAATTACTGTTCCTTGAGAAGCACACGGT |
| barcode9 | CCAGGATCCTAGGCTGCTGCTATTCCGCGGTCCGGCTAGATTGAGAAGCACACGGT |
| barcode10 | CCAGGATCCTAGGCTGCTGCATCCTACTCTTGTTACCTGATTGAGAAGCACACGGT |
| barcode11 | CCAGGATCCTAGGCTGCTGCTTGTACCTATAATCTCCTTTTTGAGAAGCACACGGT |
| barcode12 | CCAGGATCCTAGGCTGCTGCGCCCTAATGTTGCAGGGGTATTGAGAAGCACACGGT |

| | |
|---|---|
| barcode13 | CCAGGATCCTAGGCTGCTGCTCGGCTAAGTAAGAAATTCATTGAGAAGCACACGGT |
| barcode14 | CCAGGATCCTAGGCTGCTGCATGTAATGCACGAGCACCGGTTGAGAAGCACACGGT |
| barcode15 | CCAGGATCCTAGGCTGCTGCCTATATTCCTATCGTACACATTGAGAAGCACACGGT |

### *In vitro* RNA substrate

5'FAM-UCCGUUUCCUCUC**A**CCGUCGUCUCCUGCCG-3'

### Supplementary references

1. Gaudelli, N.M., Komor, A.C., Rees, H.A., Packer, M.S., Badran, A.H., Bryson, D.I., and Liu, D.R. (2017). Programmable base editing of A.T to G.C in genomic DNA without DNA cleavage. Nature *551*, 464–471. https://doi.org/10.1038/nature24644.

2. Lapinaite, A., Knott, G.J., Palumbo, C.M., Lin-Shiao, E., Richter, M.F., Zhao, K.T., Beal, P.A., Liu, D.R., and Doudna, J.A. (2020). DNA capture by a CRISPR-Cas9–guided adenine base editor. Science (1979). *369*, 566–571. https://doi.org/10.1126/science.abb1390.
